# The anti-caspase 1 inhibitor VX-765 reduces immune activation, CD4^+^ T cell depletion, viral load and total HIV-1 DNA in HIV-1 infected humanized mice

**DOI:** 10.1101/2022.09.01.502964

**Authors:** Mathieu Amand, Philipp Adams, Rafaela Schober, Gilles Iserentant, Jean-Yves Servais, Michel Moutschen, Carole Seguin-Devaux

## Abstract

**Background:** HIV-1 infection results in the activation of inflammasome involving NLRP3, IFI16, caspase-1 and release of IL-1 β and IL-18. Early inflammasome activation may facilitate viral spread and establishment of the viral reservoir. We evaluated the effect of the caspase-1 inhibitor VX-765 on virological and immunological parameters after HIV-1 infection in humanized mice.

**Methods:** NSG mice were engrafted with human CD34^+^ hematopoietic stem cells and were infected with HIV-1 JRCSF. 15 mice were first sacrificed serially to investigate kinetics of the HIV-1 related inflammasome activation. Infected mice (n=24) were then treated with VX-765 or vehicle from day 1 post infection for 21 days. Blood and organs were collected at different time points, and analysed for inflammasome genes expression, cytokines levels, viral load, CD4 cell count, and total HIV-1 DNA.

**Results:** Expression of caspase-1, NLRP3 and IL1-β was increased in lymph nodes and bone marrow on day 1 and 3 post infection (mean fold change (FC) of 2.08, 3.23, and 6.05, p< 0.001 respectively between day 1 and 3). IFI16 expression peaked at D24 in lymph node and bone marrow (FC 1.49 and 1.64, p<0.05) and coincides with increased IL-18 levels in plasma (6.89 vs. 83.19 pg/ml, p=0.004). AIM2 and IFI16 expression correlated with increased viral load in tissues (p<0.005 for the spleen) and loss of CD4^+^ T cells percentage in blood (p<0.0001 for the spleen). Treatment with VX-765 significantly reduced TNF-α at day 11 (0.47 vs. 2.2 pg/ml, p=0.045), IL-18 at day 22 (7.8 vs 23.2 pg/ml, p=0.04), CD4^+^ T cells (44.3% vs 36,7%, p=0.01) and the CD4/CD8 ratio (0.92 vs 0.67, p=0.005) in plasma. Importantly, viral load (4.26 vs. 4.89 log 10 copies/ml, p=0.027) and total HIV-1 DNA (1 054 vs. 2 889 copies /10^6^ cells, p=0.029) were decreased in VX-765-treated mice as compared to vehicle-treated mice.

**Discussion:** we report here an early inflammasome activation before detectable viral dissemination in humanized mice. We demonstrated that targeting inflammasome activation early after HIV-1 infection may represent a potential therapeutic strategy to prevent CD4^+^ T cell depletion as well as to reduce immune activation, viral load and the HIV-1 reservoir formation.

## Introduction

Around 38 million people worldwide are living with the human immunodeficiency virus (HIV) and every year, 2.1 million people get newly infected (1). Although combined antiretroviral therapy (cART) can suppress viremia and dramatically improved the life expectancy and the clinical outcome of HIV-1 infected patients (2), optimal cART treatment is not curative due to the persistence of a lifelong HIV-1 replication competent reservoir (3). In addition, cART does not fully resolve HIV-associated immune abnormalities including low grade chronic inflammation, immune activation/dysfunction and poor CD4^+^ T cell reconstitution (4, 5) and imposes a high burden to health care system due to treatment resistance, adverse events and high treatment costs. Recent evidence indicates that inflammation drives HIV-1 reservoirs persistence which in turn contributes to inflammation, creating a pathogenic vicious cycle (6). These inflammatory and immune disorders predispose HIV-1 infected patients under cART to non-AIDS age-related comorbid conditions such as cardiovascular diseases, cancer and neurodegenerative diseases and are predictive of an increased risk of morbidity and mortality (7). Strategies targeting the mechanisms of HIV-induced inflammation are now evaluated to reduce HIV-1 reservoirs, a fundamental requirement to achieve an HIV cure (8).

HIV infection triggers immune activation/dysfunction and inflammation through several mechanisms such as direct effects of viral proteins on immune cells (9-11), persistent production of type I and II interferons driven by activation of toll-like receptors with HIV-1 RNA (12), fibrosis of lymphoid structure linked to CD4^+^ T cells depletion (13) and mucosal barriers damages associated with microbial translocation (14, 15). Although inflammation can contribute to the early control of HIV infection, immune dysfunction and chronic inflammation can in turn promote viral spread and the replenishment of the reservoir by stimulating the migration of CD4^+^ T cells to sites of viral replication (16,17), generating new activated target CD4^+^ T cells (18). This leads further to reduce the ability of the immune system to kill infected cells, to increase cell-surface expression of immune checkpoints which contributes to the survival of infected cells (19, 20), and to stimulate latently infected cells to proliferate but also to produce the virus (21,22). Altogether, inflammation, poor antiviral immune response and HIV-1 persistence creates a pathogenic cycle. Interfering with mechanisms linking inflammation and HIV-1 persistence could contribute to an HIV cure and the management of inflammation-linked comorbidities. Micci et al. (23) established the proof of concept that targeting chronic inflammation with IL-21, a potent immunomodulatory cytokine, in ART-treated SIV-infected non-human primates could improve disease prognosis by reducing immune activation and the viral reservoir. To achieve HIV cure, as the viral reservoir is set within few days after infection (24), early initiation of novel treatment is required to block reservoirs establishment and may protect the immune system (25).

Inflammasomes plays important roles in innate immunity and inflammation as multiprotein signalling platforms sensing pathogens, danger signals and regulating cellular functions (26). Upon recognition of the microorganism or danger signals, certain Pattern Recognition Receptors (PRRs) recognize the presence of unique microbial components, so called pathogen-associated molecular patterns (PAMPs) or damage-associated molecular patterns (DAMPs), and recruit a multiprotein signalling platform containing the adapter protein ASC (apoptosis-associated speck-like protein containing a caspase activation and recruitment domain) which induces the proteolytic activation of the procaspase-1 (26). Subsequently, active caspase-1 initiates the release of pro-inflammatory cytokines such as IL-1β and IL-18 and the induction of pyroptosis, an highly inflammatory form of cell death. HIV-1 was described to induce the formation and action of an inflammasome in several cell types and models (27-40). Levels of IL-1β and IL-18 were found to be elevated in HIV-1 infected individuals (28-30) and to induce HIV-1 production (31-33). Ahmad et al. found that a higher proportion of blood monocytes from HIV-1 infected patients were ASC specks positive, a hallmark of inflammasome activation (34).

In innate immune cells, such as monocytes/macrophages (35,36), dendritic cells (37,38) and microglial cells (39), HIV-1 infection results in the activation of an inflammasome involving the PRR nucleotide-binding oligomerization domain (NOD), leucine-rich repeat (LRR)-containing proteins (NLR) family member 3 (NLRP3). Upon activation of NLRP3 inflammasome, pro-IL-1β is processed by activated caspase 1 to form active IL-1β. NLRP3 inflammasome activation requires a priming signal and an activation signal ultimately leading to pro-inflammatory cytokine release and pyroptosis. More specifically, in human monocytes/macrophages, HIV-1 triggers the NLRP3 inflammasome through the detection of HIV-1 RNA by TLRs and the activation of the NF-κB pathway (40). Interferon (IFN)-inducible protein 16 (IFI16) and AIM2 are DNA sensor of the AIM2-like receptors (ALRs) family and can form an inflammasome with ASC upon detection of nuclear DNA. IFI16 is also able to sense HIV-1 in macrophages, but it is unknown whether it can activate an inflammasome in such cell type (40). Using lymphoid aggregate culture (HLAC) *ex vivo* systems from fresh human tonsil or spleen tissues, Doitsch et al. revealed how most of CD4^+^ T cells dies during HIV-1 infection (41). Over 95% of CD4 T cells are non-permissive to HIV-1 infection and accumulates incomplete DNA transcripts in their cytosol. These transcripts are detected by IFI16, a sensor which activates caspase-1 resulting in the triggering of pyroptosis and the release of intracellular content and pro-inflammatory cytokines including IL-1β (42). The role of the the inflammasome in early SIV pathogenesis has recently being shown in the impairment of the innate and adaptive immune response (43) thereby facilitating viral replication (44).

Altogether, these results strongly suggest that inflammasome activation occurs early during HIV-1 infection, and could be a central mechanism in the early viral escape to the immune system and furthermore, may facilitate the establishment of the pathogenic cycle of HIV-1 reservoirs persistence and inflammation. VX-765, a selective caspase 1 inhibitor, reduces activation and activity of caspase 1 and antagonize NLRP3 inflammasome assembly and activation in the context of several inflammatory disorders or diseases such as atheroschlerosis, sepsis, or alzheimer disease (45-47). Therefore, the aim of our study was to investigate the effects of the Caspase-1 inhibitor VX-765 in an humanized mouse model of HIV-1 infection as a therapeutic strategy to reduce HIV-1 induced inflammation and reservoirs establishment.

## Results

### Early inflammasome-related genes expression upon HIV-1 infection

Humanized mice recapitulate key aspects of HIV-1 infection and pathogenesis (48, 49), however whether they recapitulate inflammasome activation upon HIV-1 infection remains undetermined. We first investigated whether inflammasome activation contributes to the early host response during HIV-1 infection in humanized mice. We generated human CD34^+^ hematopoietic stem cell-engrafted NOD/SCID/IL2rγ^null^ (huNSG) mice and performed serial necropsies on day 0 (n=6), day 1 (n=5), day 3 (n=5), day 10 (n=5) and day 24 (n=5) post intraperitoneal injection of 5×10^4^ TCID_50_ HIV-1_JRCSF_. Viremia was detectable in 3 (60%) and 5 (100%) animals sacrificed on day 10 and 24 post infection (p.i.), respectively (Supplementary Figure S1A). Acute HIV-1 infection in huNSG mice was associated with a profound decrease in CD4^+^ T cell percentages in the blood at day 24 when compared to animals sacrificed at day 10 post infection. (Supplementary Figure S1B).

Tissue viral RNA levels in hCD45^+^ cells from several tissues were further quantified by RT-PCR (Supplementary Figure S2). All huNSG mice necropsied at day 1 following HIV-1_JRCSF_ challenge were negative for viral RNA whereas HuNSG mice sacrificed at day 3 displayed at least one tissue positive for viral RNA with the spleen and the lung being positive in 80% (4 of 5) and 60% (3 of 5) animals, respectively, indicating viral dissemination at least within 72 hours. Tissue viral RNA levels became detectable in the lymph nodes (5 of 5) and the blood (4 of 5) in animal sacrificed on day 10 p.i. The number of viral RNA copies varied widely among animals sacrificed at day 7 and was elevated in most of animals sacrificed at day 24.

Previous reports have highlighted increased transcripts levels of inflammasome components upon HIV-1 infection in HIV-1 infected patients (28), SIV-infected rhesus monkeys (43) and *in vitro* models infection (41). To show evidences of early inflammasome sensing of HIV-1 in huNSG mice upon HIV-1 infection, we quantified mRNA expression of genes involved in inflammasome activation (NLRP3, NLRP1, NLRC4, AIM2, IFI16, ASC, CASP-1, IL-1β and IL-18) in hCD45^+^ cells from multiple tissues collected at necropsy. We observed a bi-phasic induction of inflammasome related-genes upon HIV-1 infection in the bone marrow, lymph nodes and spleen (Figure 1). In the bone marrow, a striking but short-lasting early upregulation of inflammasome components appeared around 3 days p.i. These included the NOD-like receptor NLRP3 (p<0.05), the adaptor ASC (p<0.01) bridging NLRs such as NLRP3 to the inflammatory Caspase-1 (CASP-1) (p<0.01) and the downstream pro-inflammatory cytokines IL-1β (p<0.01) and IL-18 (p<0.01) indicating the up-regulation of the whole NLRP3 pathway. In the lymph nodes, the induction of an early inflammasome response was evident as soon as day 1 and 3 p.i. with the upregulation of the NOD-like receptors NLRP1 (p<0.05) and NLRP3 (p<0.01), AIM2 (p<0.05), CASP-1 (p<0.01) and IL-1β (p<0.01.

**Figure 1.**
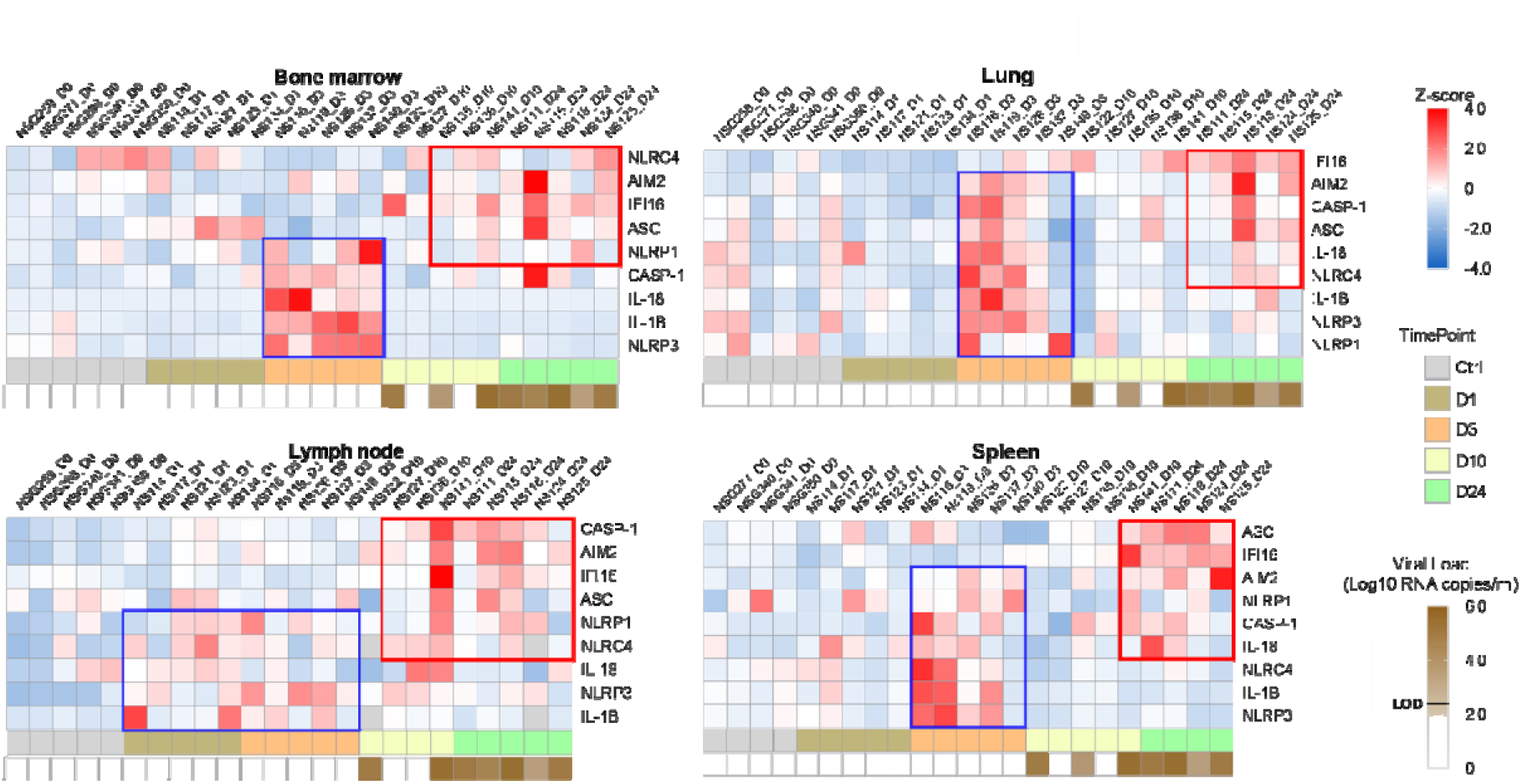
Heatmaps reveal a biphasic inflammasome-related genes transcriptomic upregulation upon HIV-1_JRCSF_ infection of huNSG mice. Relative mRNA expression of the indicated genes measured by qPCR in hCD45^+^ cells isolated from bone marrow, lung, lymph nodes and spleen in HIV-1_JRCF_ i.p. infected huNSG mice at day 0 (n=5), day 1 (n=5), day 3 (n=5), day 10 (n=6) and day 24 (n=5). Gene expression was standardized using z-score method. Red and blue colors identify up- and down-regulated genes, respectively.

Interestingly, in contrast to the bone marrow, this initial burst was sustained in the lymph nodes with the prolonged upregulation of several genes (p≤ 0.05 for NLRP3, AIM2, and CASP-1) up to day 24 p.i. In the spleen, AIM2 was the only significantly upregulated gene at 3 days p.i. (p<0.05) although we observed a tendency for other genes (NLRP3, NLRC4, CASP-1 and IL-1β.) (Supplementary Figure S3). Altogether, these rapid inflammasome-related transcriptomic changes indicate an early sensing of HIV-1 infection, especially in the bone marrow and lymph nodes, before HIV-1 RNA was quantifiable in the tissues (Supplementary Figure S2) and may reflect the host response to the inoculum challenge virus.

### IFI16 and AIM2 expression correlates with HIV-1 replication and pathogenesis

IFI16 and AIM2 are DNA sensor of the ALRs family and can form an inflammasome with ASC upon detection of nuclear DNA (50). Importantly, IFI16 restricts HIV-1 infection in macrophages but promotes HIV-1 induced CD4^+^ T cells death by pyroptosis and thus may drive HIV-1 pathogenesis (41, 42). In our humanized mice model, after the initial burst, delayed transcriptomic changes occurring around day 24 p.i. were observed, notably in the spleen. Interestingly, the upregulation of IFI16 was common in all three organs, with a maximal transcription at 24 days p.i. (p*<*0.01, 0.05 and 0.01 in bone marrow, lymph nodes, and spleen) (Supplementary Figure S3). In accordance, viremic mice demonstrated higher levels of IFI16 expression in the organs than in non-viremic mice (p<0.0001, 0.001, 0.001 in bone marrow, lymph nodes, and spleen) (Figure 2A). Furthermore, IFI16 expression in each organ correlated with corresponding tissue viral RNA copies (Figure 2B; p*<*0.0001, 0.0212 and =0.0001 in bone marrow, lymph nodes, and spleen) and with CD4^+^ T cells percentage in the blood (Figure 2C; *P*=0.0635; 0.0115 and <0.0001 in bone marrow, lymph nodes, and spleen).

**Figure 2.**
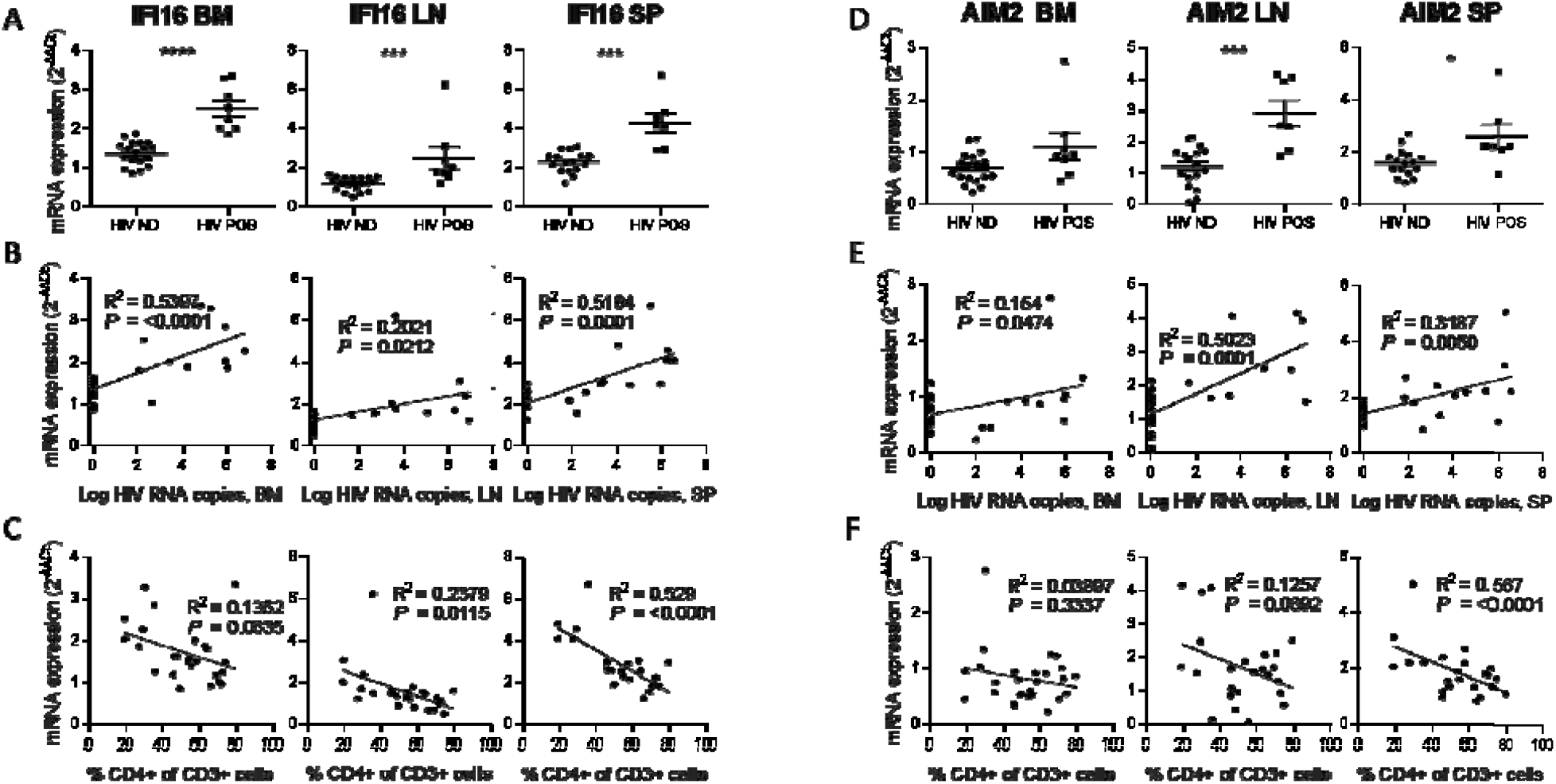
IFI16 and AIM2 expression correlates with HIV-1 replication and pathogenesis. IFI16 and AIM2 relative mRNA expression comparison between viremic (HIV POS) and aviremic (HIV ND) huNSG mice (A and D) in the bone marrow (BM), lymph nodes (LN), and the spleen (SP). Correlation between IFI16 or AIM2 relative mRNA expression with corresponding tissue viral RNA copies (B and E) or CD4 T cells percentage in the blood (C and F). qPCR results were analyzed using the 2^-ΔΔCT^ method. Statistical tests were analysed by Mann–Whitney t-tests for comparison of two groups or Spearman correlation tests.

Similarly, AIM2 reached a maximum of expression at day 24 p.i. in the lymph nodes and the spleen (p<0.01 and 0.01, respectively) (Supplementary Figure S3) and was expressed at higher levels in viremic mice when compared to non-viremic mice (p<0.001 and 0.05 in lymph nodes and spleen) (Figure 2D). AIM2 expression also correlated with viral RNA copies in the same organ (Figure 2E; p=0.0474, 0.0001 and 0.005, in bone marrow, lymph nodes and spleen) and inversely with CD4^+^ T cells percentage in the blood (Figure 2F; p<0.0001 for AIM2 in the spleen). In addition, other inflammasome related genes (ASC and CASP-1) reached a maximum of expression 24 days p.i., notably in the spleen (*P*<0.01; ASC, CASP-1) and in the lymph nodes (*P*<0.01; CASP1) (Supplementary Figure S3) and their expression was associated with HIV-1 replication and pathogenesis (Supplementary Figure S4).

### HIV-1 infection induces the release of inflammasome-related cytokines

Typical inflammasome activation requires a two-step process including (1) a priming signal resulting in the transcriptional upregulation of inflammasome-related genes and (2) a second signal causing inflammasome assembly and activation at the protein level. Upon assembly, the inflammasome induces the proteolytic activation of procaspase-1 into active caspase-1 that initiates the maturation and release of inflammasome-related pro-inflammatory cytokines such as IL-1β and IL-18 (50). Therefore, qPCR measurements of inflammasome-related gene expression mostly reflects the priming steps (Figure 2 and Supplementary Figure S3) while inflammasome activation usually requires the quantification of mature IL-1β and IL-18 or caspase-1 activity.

To confirm inflammasome activation in huNSG mice upon HIV-1 infection, we next quantified plasmatic levels of inflammasome-related (IL-1β and IL-18) and others pro-inflammatory cytokines (IFNγ, IL-6, IP-10 and TNFα). Plasma was collected at necropsy on day 0 (n=5), day 3 (n=5), day 10 (n=5) and day 24 (n=6) post infection. HIV-1 infection was previously described to induce a rapid and intense cytokine storm involving several pro-inflammatory biomarkers (51). In accordance, we observed a rapid elevation in the plasmatic levels of several pro-inflammatory cytokines such as IFNγ, IL-18, IP-10 and TNFα upon HIV-1 infection (Figure 3A and Suppl Figure 5). Interestingly, IL-18 levels were increased at day 10 and day 24 post infection (Figure 3A; *P* <0.05 and 0.01, respectively) and viremic mice harboured higher IL-18 levels than non-viremic mice (Figure 3B; *P*<0.001). Furthermore, its plasmatic levels correlated with viremia (Figure 3C; *P*=0.0548) and inversely with CD4^+^ T cells percentage in the blood (Figure 3D; p=0.0212). In addition to IL-18, IL-1β levels tended to increase at day 3 post infection (Figure 3A) but this was not significant. Its short half-life (52), low volume of plasma collected in mice, and plasmatic levels below the LOD in several HuNSG mice might explain that we did not catch a significant early increase for this cytokine. Altogether, these results strongly indicates that an inflammasome is activated upon HIV-1 infection in our humanized mice model and is associated with disease progression.

**Figure 3.**
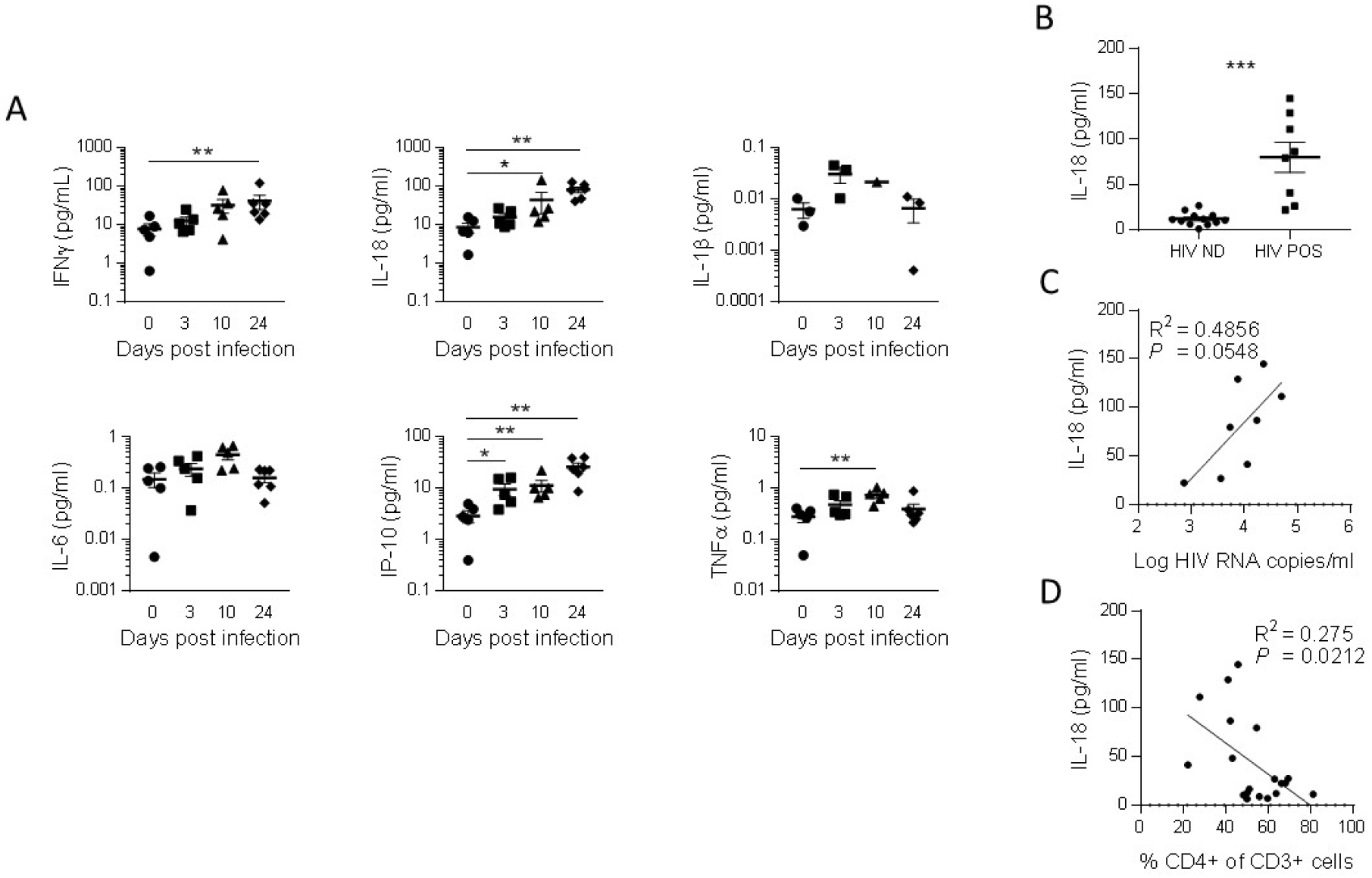
IL-18 is induced by HIV-1 replication and correlates inversely with CD4^+^ T cells percentage. (A) Plasmatic cytokines concentration measured using a multiplex U-PLEX Biomarker assay kit in the plasma of HIV-1_JRCF_ i.p. infected huNSG mice. (B) Plasmatic IL-18 levels comparison between viremic (HIV POS) and aviremic (HIV ND) huNSG mice. Correlations between plasmatic IL-18 levels with the plasmatic viral load (C) or CD4^**+**^ T cells percentage in the blood (D) at day 0 (n=5), day 3 (n=5), day 10 (n=6) and day 24 (n=5). Statistical tests were analysed by Mann– Whitney t-tests for comparison of two groups or Spearman correlation tests.

### Inflammasome inhibition reduces HIV-1 induced circulating levels of IL-18 and the cytokine storm in HIV-1 infected HuNSG mice

To decipher the impact of inflamasomme on HIV-1 pathogenesis *in vivo*, we treated HuNSG mice daily by intraperitoneal injection of the caspase-1 inhibitor VX-765 (53) at the dose of 200 mg/kg (54, 55) for 21 consecutive days starting on day 2 after HIV-1 infection to cover the early upregulation of inflammasome-related genes. As expected, HIV-1 infection in the vehicle group induced a potent cytokine storm with elevated plasmatic levels of the inflammasome-related cytokines IL-1β and IL-18 but also IFNγ, IL-6, IP-10 and TNFα when compared to day 10 before infection (*P*<0.05; 0.01; 0.05; 0.001; 0.01; 0.001, respectively; Supplementary Fig. 6). In contrast, caspase-1 inhibition blunted the amplitude of the HIV-1 induced cytokine storm as VX-765 treated HuNSG mice only upregulated significantly the levels of IP-10 and TNFα after infection (*P*<0.05 and 0.05, respectively; Supplementary Fig. 6E and F). Importantly, caspase-1 inhibition reduced the circulating levels of the inflammasome-related cytokine IL-18 at day 22 in comparison with the vehicle group (*P*<0.05; Figure 4B). TNFα levels at day 10 were also reduced under VX-765 treatment (*P*<0.05, Figure 4F) when compared to the vehicle group. Although IL-1β levels increased at day 10 post infection in the vehicle group when compared to day 10 before infection, differences between VX-765 and vehicle group were not significant (Figure 4C), probably due to low concentration measured at this time point. Altogether, these results confirm that the inhibition of inflammasome upon HIV-1 infection in HuNSG mice reduced the levels of circulating pro-inflammatory cytokines induced during early HIV-1 infection.

**Figure 4.**
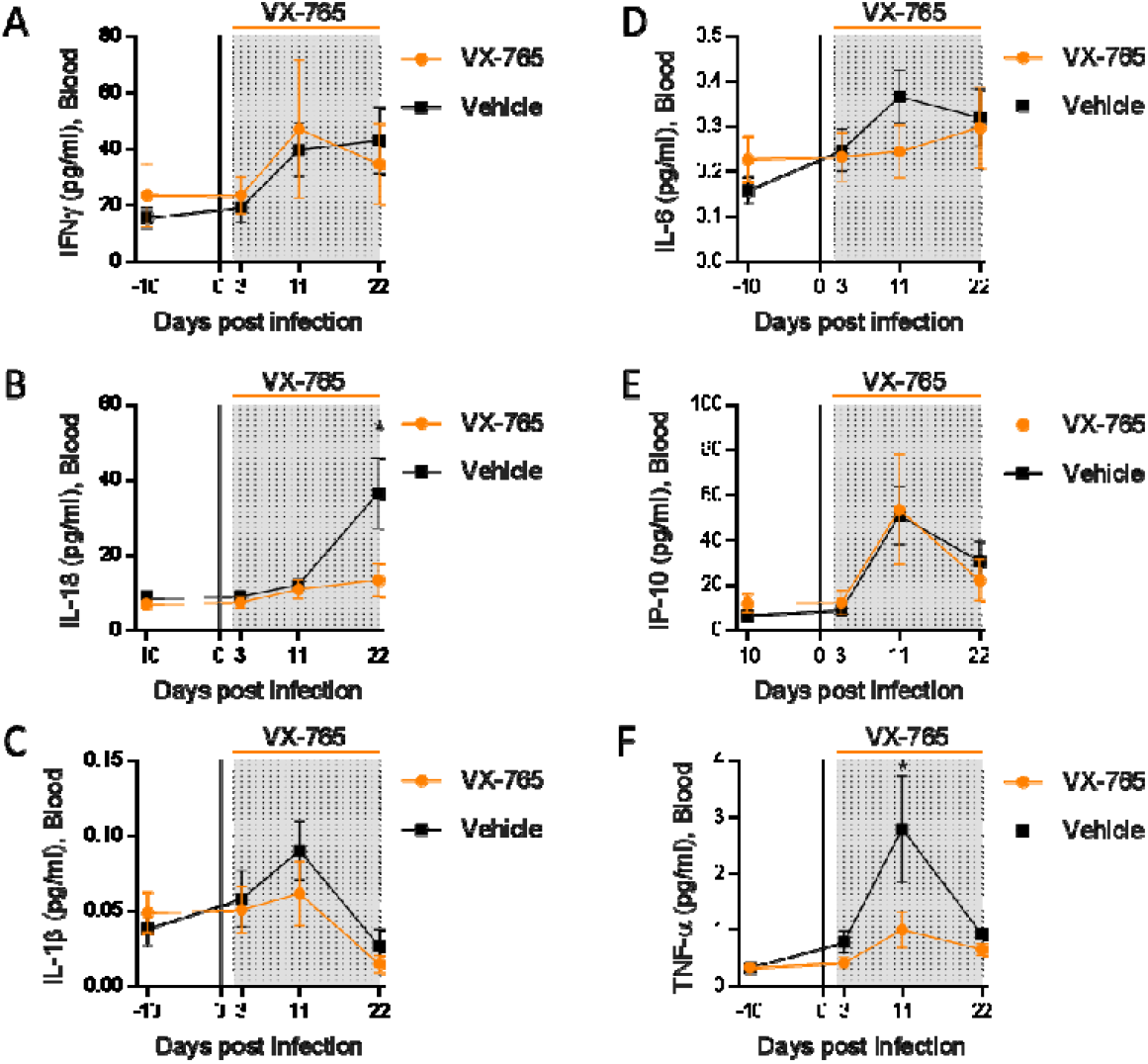
Caspase-1 inhibition reduces circulating levels of IL-18. Plasmatic cytokines concentration measured using a multiplex U-PLEX Biomarker assay kit in longitudinal plasma samples of HIV-1_JRCF_ i.p. infected huNSG mice treated with VX-765 (VX-765, n=12) or vehicle (Vehicle, n=12). Statistical tests were analysed by Mann–Whitney t-tests for comparison of two groups.

### Administration of VX-765 prevents CD4^+^ T cell depletion and decreases viral load and total HIV DNA in HIV-1 infected HuNSG mice

Inflammasome activation have been proposed as the main mechanism of CD4^+^ T cells death during HIV infection (41, 42). Therefore, we monitored CD4^+^ T cells percentages and CD4/CD8 T cell ratio longitudinally in the blood and at necropsy in the organs after the administration of the caspase-1 inhibitor. Vehicle and VX-765 treated groups presented a significant redution of CD4^+^ T cells percentages and CD4/CD8 T cell ratio in the blood at day 22 post infection (Suppl Fig. 7A and B). Interestingly, althought CD4^+^ T cells percentages and CD4/CD8 T cell ratio were equivalent in both groups before HIV-1 infection and VX-765 treatment, percentages of circulating CD4^+^ T cells and CD4/CD8 T cell ratio at day 22 post infection in the VX-765 treated group were slightly but significantly higher (p<0.05; Figure 5A and C). At necropsy, we found that CD4^+^ T cells percentages and CD4/CD8 T cell ratio in the bone marrow but not in the sleen were also higher in the VX-765 treated HIV-1 infected HuNSG compared to the vehicle group (p<0.05 and n.s.; Figure 5B, C, E, F).

**Figure 5.**
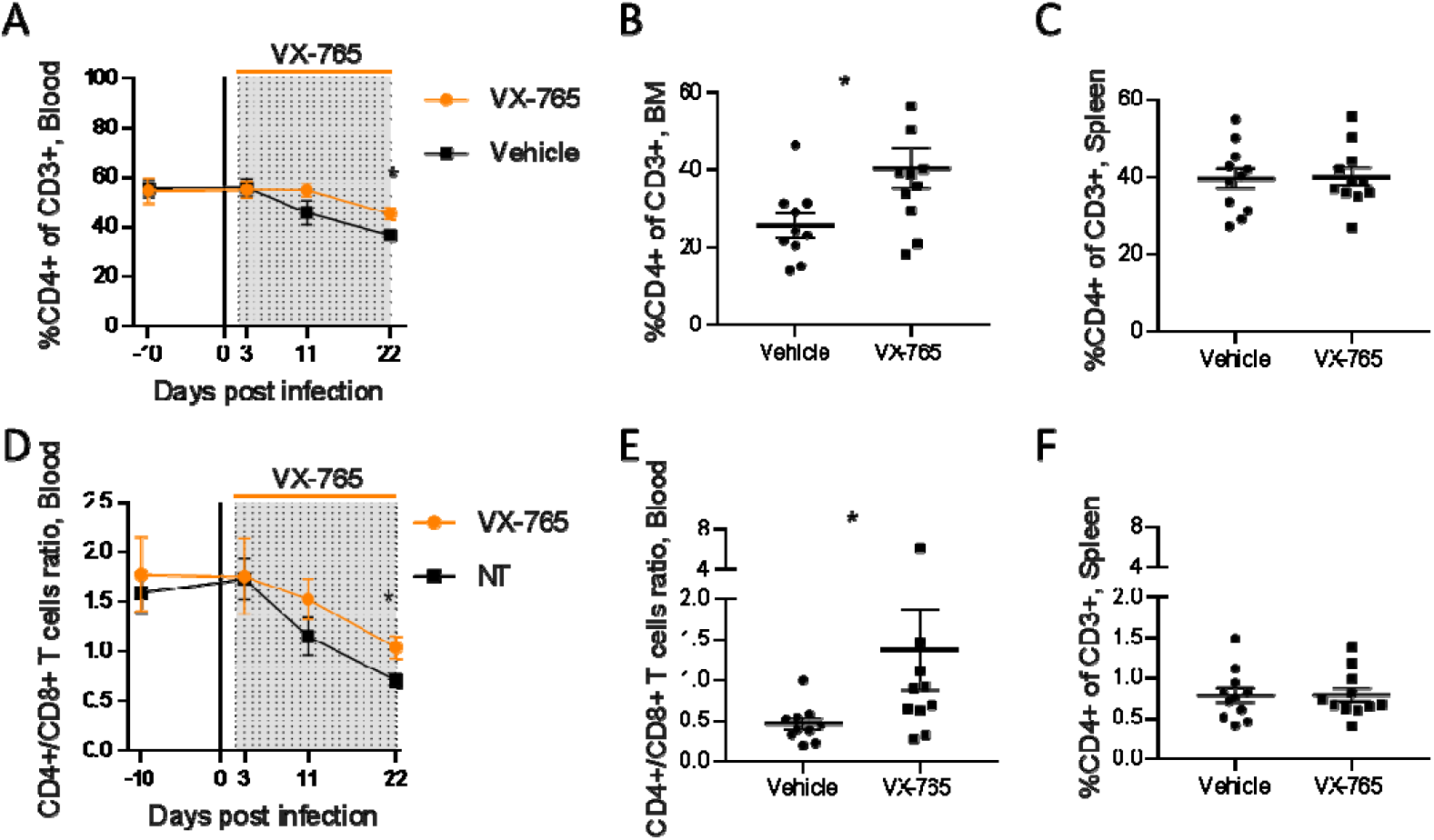
VX-765 treatment prevents CD4^+^ T cell depletion in the blood and bone marrow (BM) during HIV-1 infection of huNSG mice. Blood human CD4^+^ T cells percentage among CD3^+^ cells (A) and CD4^+^/CD8^+^ T cell ratios (D) measured longitudinally by flow cytometry in vehicle or VX-765 treated HIV-1 infected HuNSG mice. Bone marrow (BM) (B and E) and spleen (C and F) human CD4^+^ T cells percentage among CD3^+^ cells and CD4^+^/CD8^+^ T cell ratios, respectively, measured at day 22 post infection in vehicle or VX-765 treated HIV-1 infected HuNSG mice. Statistical tests were analysed by Mann–Whitney t-tests for comparison of two groups.

In addition to reducing CD4^+^ T cell depletion, we observed lower plasmatic viral loads in VX-765 treated mice (p<0.05; Figure 6A, 6B, 6C) indicating that inflammasome inhibition interferes with HIV-1 replication. When measuring total HIV-1 DNA in human CD45^+^ cells from the spleen of all animals, we also found a reduced content of total HIV-1 DNA (Figure 6D) in VX-765 treated mice as compared to the vehicle treated mice (1 054 vs. 2 889 copies / 10^6^ cells, p=0.029).

**Figure 6.**
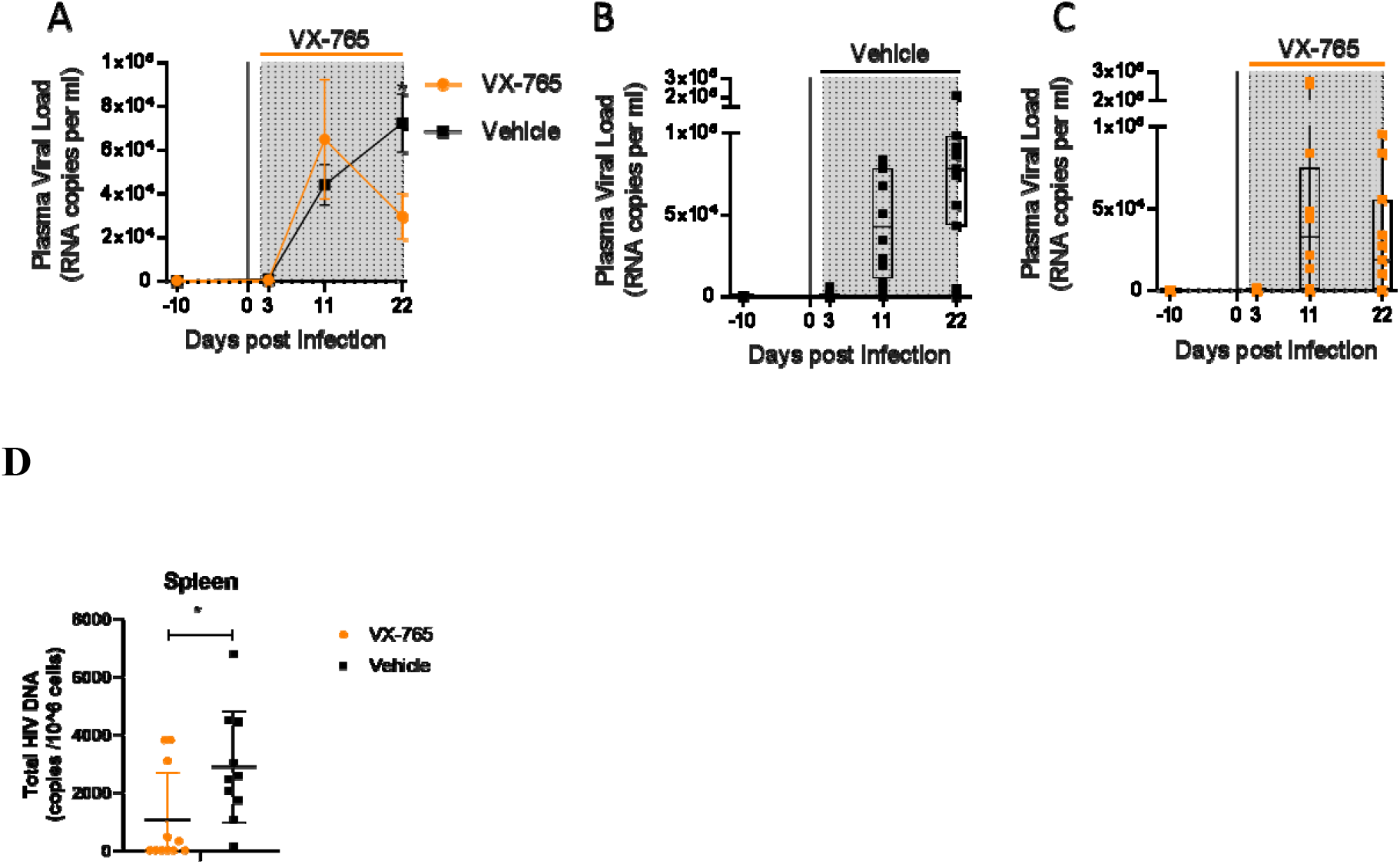
VX-765 treatment reduces plasma viral load and total HIV-1 DNA during HIV-1 infection of huNSG mice. Plasma viral load (A) was measured longitudinally by ddPCR in vehicle (B) or VX-765 (C) treated HIV-1 infected HuNSG mice. Total HIV-1 DNA was measured by qPCR in the spleen of CD45^+^ human cells isolated by MACS purification from all animals treated or not with VX-765 (D). Statistical tests were analysed by Mann–Whitney t-tests for comparison of two groups.

### VX-765 inhibits *in vivo* caspase-1 activity in splenic CD11c+ and CD14+ cells

To better understand the effect of VX-765 treatment on inflammasome activation in immune cells subsets, we next investigated caspase-1 activity using the FAM-FLICA® Caspase-1 assay reagent at day 23 post infection in CD4^+^, CD8^+^, CD14^+^ and CD11c^+^ cells from the spleen and the bone marrow in VX-765 and vehicle treated HIV-1 infected HuNSG mice (Supplementary Fig. 8). In this assay, the fluorescent FAM-YVAD-FMK probe binds irreversibly to active caspase-1 in living cells. We found the highest caspase-1 activity among CD11c^+^ and CD14^+^ cells (Supplementary Fig. 8 and Figure 7).

**Figure 7.**
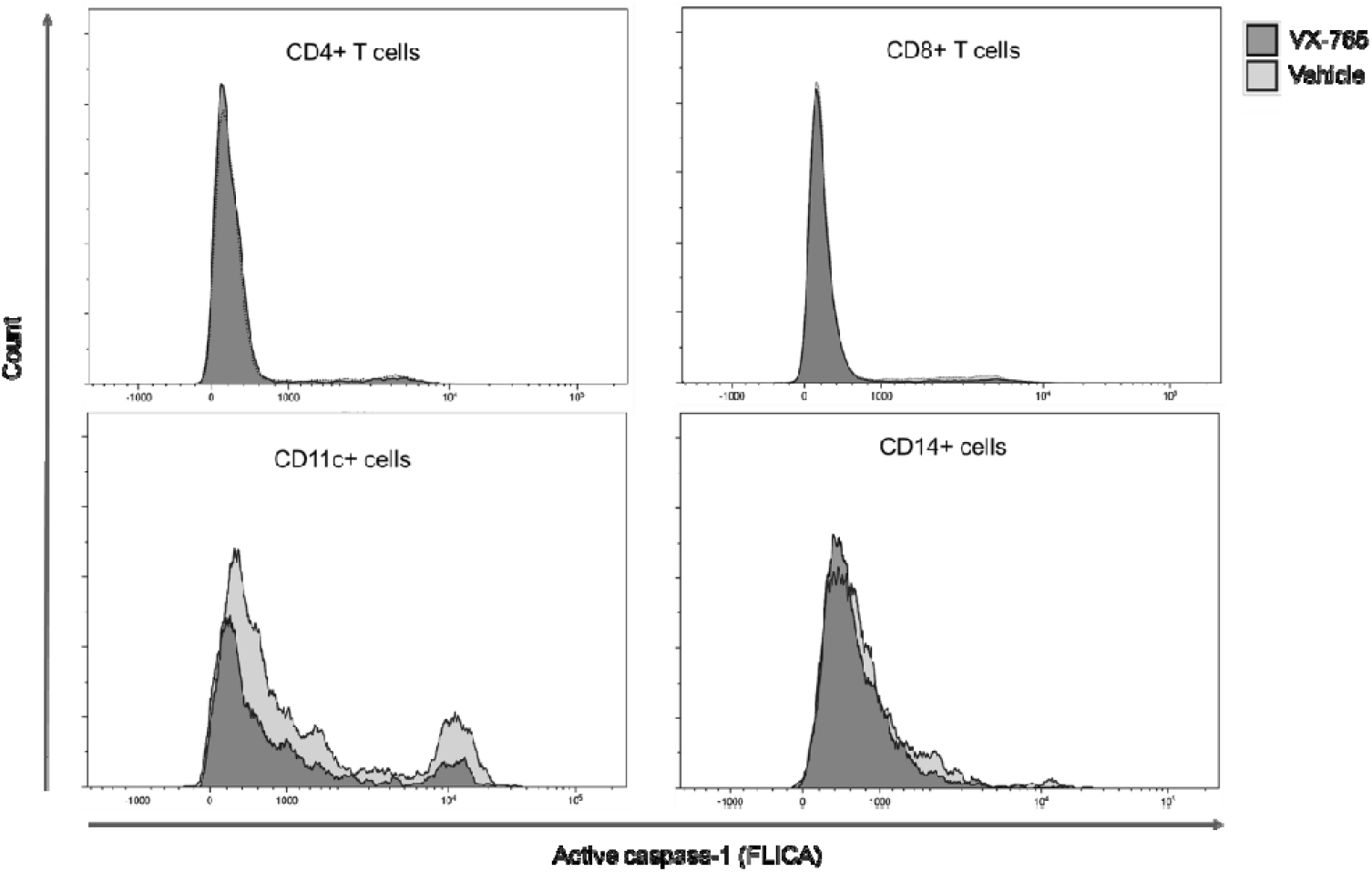
Representative histograms of FLICA staining in splenocytes from VX-765 and vehicle treated HIV-1 infected HuNSG mice. Splenocytes were stained for active caspase-1 using FAM-FLICA reagent. Representative MFI of FAM-FLICA^+^ CD4^+^, CD8^+^, CD11c^+^ and CD14^+^ splenic cells in one HIV-1 infected HuNSG mice treated with vehicle or with VX-765 treated mice at day 22 post-infection is shown.

Importantly, splenic CD11c^+^ and CD14^+^ cells presented a reduced caspase-1 activity upon VX-765 treatment (Figure 8A, B, C, D). On the contrary, CD4^+^ and CD8^+^ T cells presented low levels of caspase-1 activity and did not respond to VX-765 treatment (Figure 9A, B, C, D). In the bone marrow, caspase-1 activity was lower in all subsets compared to the spleen, and none of them presented a decreased caspase-1 activity in response to VX-765 treatment (Supplementary Fig.9). These results indicates that inflammasome activation *in vivo* rather takes places in splenic innate immune CD11c^+^ and CD14^+^ cells than splenic adaptive CD4^+^ or CD8^+^ or bone marrow immune cells. Furthermore, we found a considerable positive correlation between the percentage of active caspase-1 splenic CD11c^+^ cells from the vehicle treated group and the circulating levels of IL-18 (P<0.0001; Figure 8E). Such correlations were not significant in other subsets (Figure 8F and Figure 9E and 9F) and indicates that HIV-1 induced inflammasome activation in splenic CD11c^+^ cells may be the principal source of circulating IL-18.

**Figure 8.**
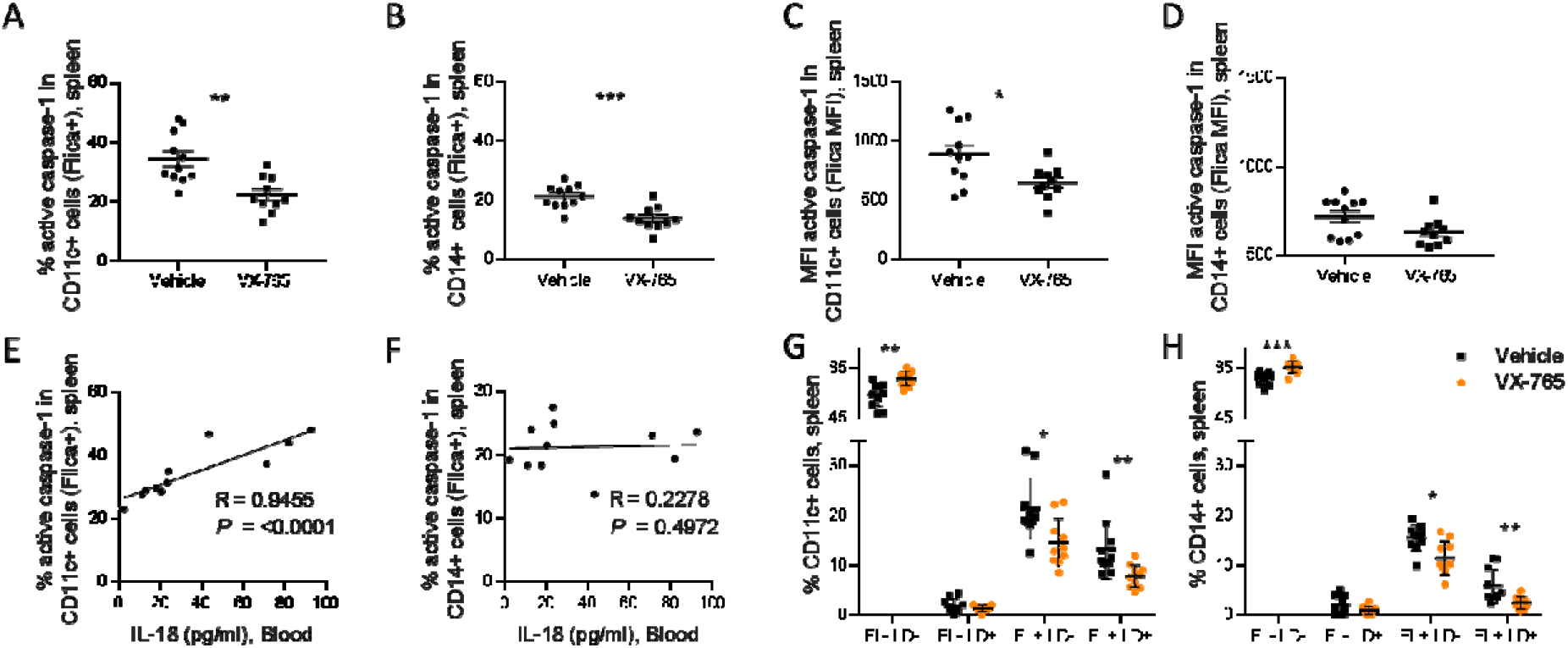
VX-765 treatment reduces caspase-1 activation in splenic CD11c^+^ and CD14^+^ in HIV-1 infected HuNSG mice. Splenocytes were stained for active caspase-1 using FAM-FLICA reagent. Percentage and MFI of FAM-FLICA^+^ CD11c^+^ (A, C), CD14^+^ (B, D) splenic cells in vehicle or VX-765 treated HIV-1 infected HuNSG mice at day 22 post-infection. Correlation between the percentage of FAM-FLICA^+^ CD11c^+^ (E), CD14^+^ (F) splenic cells and IL-18 plasmatic levels in vehicle treated HIV-1 infect group at day 22 post-infection. Percentage of FAM-FLICA^-^/Live-dead (FL^-^LD^-^), FAM-FLICA^-^/Live-dead^+^ (FL^-^LD^+^), FAM-FLICA^+^/Live-dead^-^ (FL^+^LD^-^) and FAM-FLICA^+^/Live-dead^+^ (FL^+^LD^+^) CD11c^+^ (G), CD14^+^ (H) splenic cells in vehicle or VX-765 treated HIV-1 infected HuNSG mice at day 22 post-infection. Statistical tests were analysed by Mann–Whitney t-tests for comparison of two groups or Spearman correlation tests.

**Figure 9.**
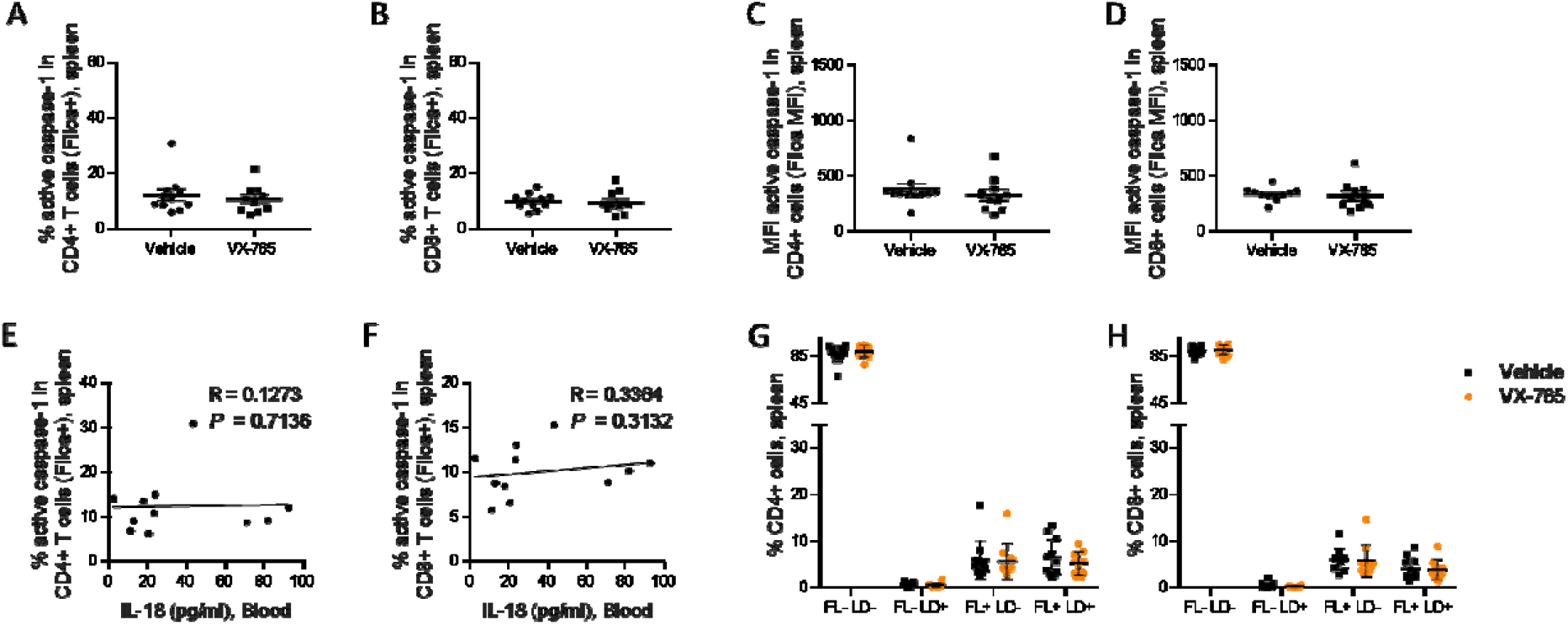
VX-765 treatment does not affect caspase-1 activation in splenic CD4^+^ and CD8^+^ T cells in HIV-1 infected HuNSG mice. Splenocytes were stained for active caspase-1 using FAM-FLICA reagent. Percentage and MFI of FAM-FLICA^+^ CD4^+^ (A, C), CD8^+^ (B, D) splenic cells in vehicle or VX-765 treated HIV-1 infected HuNSG mice at day 22 post-infection. Correlation between the percentage of FAM-FLICA^+^ CD4^+^ (E), CD8^+^ (F) splenic cells and IL-18 plasmatic levels in vehicle treated HIV-1 infect group at day 22 post-infection. Percentage of FAM^-^FLICA^-^/Live-dead^-^ (FL^-^LD^-^), FAM-FLICA^-^/Live-dead^+^ (FL^-^LD^+^), FAM-FLICA^+^/Live-dead^-^ (FL^+^LD^-^) and FAM-FLICA^+^/Live-dead^+^ (FL^+^LD^+^) CD4^+^ (G), CD8^+^ (H) splenic cells in vehicle or VX-765 treated HIV-1 infected HuNSG mice at day 22 post-infection. Statistical tests were analysed by Mann–Whitney t-tests for comparison of two groups or Spearman correlation tests.

In addition to the release of pro-inflammatory cytokines such as IL-1β and IL-18, active caspase-1 initiates the induction of pyroptosis, an highly inflammatory form of cell death (42). To determine the effect of inflammasome inhibition on immune cells viability, a fluorescent probe (LIVE/DEAD) was used to identify dying cells in addition to caspase-1 activity. The active caspase-1 (FLICA) fluorescence was plotted against LIVE/DEAD fluorescence to obtain four quadrants: live cells negative for caspase-1 activity and LIVE/DEAD (FL^-^LD^-^), live cells positive for caspase-1 activity (FL^+^LD^-^), necrotic/late apoptotic cell (FL^-^LD^+^) and pyroptotic cells (FL^+^LD^+^) (Supplementary Fig. 10). Using theses gates, we observed that VX-765 treatment reduced both the percentages of live cells positive for caspase-1 activity and pyroptotic cells in CD11c^+^ and CD14^+^ when compared to the vehicle treated group while the percentage of necrotic/late apoptotic cells remained unchanged (Figure 8G and H). In contrast, no changes upon treatment were observed in CD4^+^ and CD8^+^ T cells (Figure 9G and H). Given the low levels of caspase-1 activity in CD4^+^ and CD8^+^ T cells together with the observation that caspase-1 activity in these cells was not inhibited using VX-765, a specific caspase-1 inhibitor, we cannot rule out the possibility that FLICA staining in these cells corresponds to a basal background rather than inflammasome activation.

## Material and methods

### Humanized mice and HIV infection

NOD.Cg-*Prkdc*^*scid*^ *Il2rg*^*tm1Wjl*^/SzJ (NSG) mice were purchased from the Jackson Laboratory. All mice were bred and maintained in microisolator cages at the specific pathogen free (SPF) animal facility of the Luxembourg Institute of Health according to national and EU regulations. All experiments on animals were performed with the authorizations from the animal welfare committee of the Luxembourg Institute of Health and the Ministry of Veterinary and Agriculture of Luxembourg. 3- to 4-weeks-old mice juvenile NSG mice were conditioned and humanized as previously described (49). Briefly, NSG mice were conditionned with two intraperitoneal injections of Busulfan (2×20 mg/kg, Busilvex®) with a 12-hr interval. Conditionned NSG mice were humanized with CD34^+^ hematopoietic stem cells isolated from human cord blood (CB) using a magnetic activated cell sorting CD34^+^ progenitor cell isolation kit (Stem Cell Technologies). CB was provided by the Cord Blood Bank Central Hospital University (Liège, Belgium) and was collected after obtaining written infomed consent. 2×10^5^ freshly isolated CD34^+^ cells were transplanted intravenously (i.v.). Twenty four weeks post-transplantation, humanized mice with more than 10% human CD45^+^ cells in the peripheral blood were transferred to level 3 animal facilities and infected by intraperitoneal injection of a HIV laboratory adapted strain (JRCSF, 10.000 TCID50).

### VX-765 treatment

VX-765 (Invivogen) was dissolved in 20% cremophor and injected intraperitoneally (i.p.) in HIV-1 infected humanized mice at 200 mg/kg once a day starting at day 2 post-infection and for 21 consecutive days. Control humanized mice received the corresponding vehicle.

### Sampling collection and cell isolation

Whole blood was collected into EDTA-coated tubes (BD Microtainer K2E tubes; BD Biosciences) from the temporal vein for longitudinal sampling or by cardiac puncture at necropsy. Plasma was separated from cells by centrifugation 10 min at 2000 rpm at 4°, aliquoted in sterile Eppendorf tubes and stored at -80°C until subsequent analysis. Peripheral blood cells were further stained for flow cytometry. Organs were collected at necropsy, washed in cold RPMI and processed immediately for cell isolation. Single cell suspensions were prepared from spleens, lymph nodes and bone marrows. Spleens and lymph nodes were gently disrupted in cold PBS using the plunger end of a syringe. Bone marrow cells were obtained from tibias and femurs. Both ends of the bones were cut and the bone marrow were flushed using a 25-gauge needle and 1 mL syringe filled with cold PBS. Cells from bone marrow and splenocytes suspensions were passed through 70 μm nylon cell strainers (BD biosciences) and centrifuged 10 min at 1200 rpm at 4°C. Single cell suspensions were further used for hCD45^+^ cells isolation, stained for flow cytometry or pelleted prior to DNA extraction.

### Human CD45^+^ cells isolation, RNA extraction and reverse transcription polymerase chain reaction

Human CD45^+^ cells were isolated from peripheral blood or spleen, bone marrow and lymph nodes single cell suspensions using CD45 MicroBeads, human positive selection kit (Miltenyi Biotec) and total RNA was extracted from hCD45^+^ cells using NucleoSpin® RNA kit (Macherey-Nagel) following manufacturer’s instructions. 200 ng of RNA was reverse transcribed into cDNA using High-Capacity RNA-to-cDNA™ Kit (Applied Biosystems). The cDNA was used for the quantification of human NLRP1 (TaqMan Assay: Hs00248187_m1), NLRP3 (TaqMan Assay: Hs00918082_m1), NLRC4 (TaqMan Assay: Hs00892666_m1), AIM-2 (TaqMan Assay: Hs00915710_m1), IFI16 (TaqMan Assay: Hs00986757_m1), ASC (TaqMan Assay: Hs00203118_m1), CASP-1 (TaqMan Assay: Hs00354836_m1), IL-1β (TaqMan Assay: Hs01555410_m1), and IL-18 (TaqMan Assay: Hs01038788_m1) genes expression by real-time PCR. ACTB gene (TaqMan Assay: Actb-Hs00357333_g1) was used as a control for expression levels normalization. PCR amplification of cDNA was done in 20 μL volume containing 10 μL of TaqMan® Fast Advanced Master Mix (2X) (Applied Biosystems), 1 μL of TaqMan® Assay (20X) (Applied Biosystems), 7 μL of Nuclease-Free Water and 2 μL of cDNA. All quantitative PCR were performed on a 7500 Real-Time PCR System (Applied Biosystem). The results were analyzed using the 2^-ΔΔCT^ method.

### Flow cytometry

Human hematopoietic cell engraftment and immunophenotyping in mouse xenorepicients was performed by flow cytometry on peripheral blood or spleen and bone marrow single cell suspensions at various timepoints. Red blood cells from peripheral blood and spleen samples were lysed using RBC lysis buffer (BD). Cells were stained for 30 min at 4°C in 5ml round bottom polypropylene tubes with fluorochrome-labeled monoclonal antibodies. The humanisation antibody cocktail contained CD4-BUV395 (SK3), CD3-BUV496 (UCHT1), CD8-BUV805 (SK1), hCD45-BV421 (HI30), mCD45-PE Cy5 (30-F11), CD14-APC (M5E2) from BD Bioscience and CD19-PE (SJ25C1) from Biolegend. For phenotyping studies CD152-BV786 (BNI3), CD69-PE CF594 (FN50) from BD and HLA DR-BV711 (L243), CD38-PerCP Cy5.5 (HIT2), CD279-PE Cy7 (EH12.2H7) from Biolegend were added to the cocktail. For the FAM-FLICA® Caspase-1 assay (ImmunoChemestry Technologies), 30X FAM-FLICA fluorescent dye was added on cells at a final ratio of 1:60 and incubated 1h at 37°C and washed with 2 ml of 1X Apoptosis wash buffer. For this experiment CD11c-PE (B-ly6) (BD), CD163-PerCP Cy5.5 (GHI/61) and CD16-PE Cy7 (3G8) (Biolegend) were added for flow cytometry staining. For all experiments, cell viability was assessed using LIVE/DEAD Near-IR dead cell stain kit from Invitrogen according to the manufacturer instructions. Cells were fixed for 1h at 4°C using 1X BD Lysing solution (Cat n°: 349202) before acquisition. Samples were acquired on a FACS Fortessa SORP 5 laser instrument (BD Biosciences) and analyzed with the Kaluza Flow Cytometry Analysis Software (Beckman Coulter).

### Total HIV-1 DNA and RNA quantification

Total DNA was extracted from spleen, lymph nodes and bone marrow cells by manual extraction with the NucleoSpin® Tissue kit (Macherey-Nagel) according to manufacturer’s instructions. Aliquots of eluted samples were frozen at −80□°C until total HIV-1 DNA quantification by ddPCR as previously described (49). Plasma viral load was measured by digital droplet PCR (ddPCR) assay as previously described (49). Thirty μL of humanized mice plasma was diluted with 70 μL of PBS and processed. The limit of detection in this assay is 235 copies/ml.

### Cytokine levels

Levels of human cytokines IL-1β, IL-6, IL-18, IFN-γ, IP-10 and TNF-α were quantified in 25 μL of plasma using a multiplex U-PLEX Biomarker assay kit (Meso Scale Diagnostics) according to the manufacturer’s instructions. Data were acquired and analysed using MESO QuickPlex SQ 120 instrument.

### Statistical analysis

Statistical analysis was performed using GRAPHPAD PRISM software. Data were expressed as the mean value ± SEM. Groups were compared using unpaired Mann-Whitney *U* test or Wilcoxon’s matched pairs signed rank test. A P-value < 0,05 was considered to be significant. Two-tailed Spearman correlation coefficient (r) was calculated, with resulting P-values at 95% confidence

## Discussion

In the current work, we showed that targeting inflammasome activation early after HIV infection using the caspase-1 inhibitor VX-765 may represent a potential therapeutic strategy to improve CD4^+^ T cell homeostasis, reduce viral load and immune activation as well as to decrease the HIV-1 reservoir. Many studies have outlined the implications of inflammasome activation during the acute and chronic phase of HIV-1 infection as recently reviewed by Leal et al (56). The activation of inflammasomes in monocytes, macrophages and dendritic cells leads to the release of cytokines and plays an important part in the first line of host defense against HIV-1 (56). However it can also promote viral spread by the recruitment and activation of CD4^+^ T cells (16, 17). Furthermore, the chronic activation of inflammasomes might be a driver of viral reservoir persistence (18), which in turn can trigger inflammation, creating a pathogenic cycle. Breaking out of this pathogenic cycle could represent an important building block to reduce non-AIDS related morbidities caused by persisting inflammation during cART (57). To date, despite increased therapeutic interests (45-47), no compound targeting the inflammasome was marketed (58). Nevertheless, The caspase 1 inhibitor VX-765 was approved by the Food and Drug Administration for human clinical trials, showed a good safety profile, and represents currently a promising drug to prevent the onset of inflammation in several diseases (47).

In this regard, we investigated in a first step whether the humanized mouse model might be a suitable pre-clinical model to evaluate anti-inflammasone therapy through the detailed kinetics of inflammasome activation in tissues from humanized mice. During the early host response to HIV-1 infection in humanized NSG mice, we described a clear bi-phasic activation profile of inflammasome related genes in bone marrow, lungs, lymph node and spleen. The rapid inflammasome-related transcriptomic changes indicate an early sensing of HIV-1 infection, before HIV-1 RNA was quantifiable in the tissues and may reflect the host response to the virus challenge. These findings are in agreement with previous studies from HIV-1 infection in humans (39) and SIV infection in rhesus monkeys (43). Interestingly, in the first phase (up to day 3 post infection), the expression of NLRP3, IL-1β and IL-18 was significantly upregulated in several organs before viremia was detectable in blood. Of note, in lymph nodes and bone marrow transcriptomic changes occurred even before detectability of the virus in the respective tissues. The genes induced in the bone marrow included NLRP3, able to respond to a diverse set of stimuli including HIV ssRNA and proteins, the adaptor ASC, bridging NLRs such as NLRP3 to the inflammatory Caspase-1, CASP-1 and the downstream pro-inflammatory cytokines IL-1β and IL-18 indicating the up-regulation of the entire NLRP3 pathway. In the lymp nodes, the induction of an early inflammasome from day 1 and day 3 p.i. on, was evident from the upregulation of the NOD-like receptors NLRP1, NLRP3, AIM2, CASP-1 and IL-1 β, indicating an early host response to HIV-1 infection in humanized mice, in agreement with results obtained from the SIV model (43, 44). After the initial burst we observed a second phase of transcriptomic changes dominated by IFI16 and AIM2 in bone marrow, lymph nodes and spleen. In response to DNA, IFI16 is able to mediate the induction of interferon-β through the STING pathway (59) but it can also form an inflammasome with ASC upon detection of nuclear DNA (60). AIM2 is a cytoplasmic double stranded DNA sensor and can initiate the assembly of the inflammasome (61-63). Previous reports indicated that IFI16 restricts HIV-1 infection trough STING in macrophages (40) but promotes HIV-1 induced CD4^+^ T cells death by pyroptosis and thus may drive HIV-1 pathogenesis (41, 42). Here we show convincing direct correlations between IFI16 but also AIM2 expression with viral load in all tissues and the inverse correlations with the percentage of CD4^+^ T cells (Figure 2). Only in the bone marrow, the AIM2 correlation did not reach statistical significance, probably due to the low mRNA quantity detected here. These results are reinforced by the similar good correlation obtained between ASC and CASP-1 expression and the percentage of CD4^+^ T cells or viral load in lymph nodes and in the spleen 24 days post-infection (Supp Figure 4). Taken together these data emphasize the dual role of inflammasome at the onset of HIV-1 infection that fight first the virus and then fuels disease progression.

Inflammasome activation by NLRP3 is a tightly regulated process requiring signals in two main steps. First a priming signal which leads to transcription of inflammasome related genes and secondly a signal initiating the assembly of the multiprotein complex. As a result, caspase-1 activity promotes the proteolytic cleavage of the pro-inflammatory cytokines IL-1β and IL-18. We report here significant increases of circulating protein levels of IL-18 at day 10 and 24 post HIV-1 infection (Figure 3) indicating a full blown inflammasome activity in humanized mice. We found also a high systemic burst of several plasma cytokines (IFN-γ, TNF-α, IP-10 and IL-18) in agreement with the charateristic cytokine storm of acute HIV-1 infection in humans. For IL-1β, we did not find a significant increase of the cytokine in the plasma of HIV-1 infected mice as compared to non-infected mice, only a weak tendancy of increased levels, in contrast to mRNA expression in all tissues. This discrepancy might be due to the short half life of the cytokine (51) and the low basal secretion of protein expression in plasma of the mice. In our humanized mice model, we used an improved engraftment protocol with busulfan myeloablation to reach T cell counts sufficient to sustain long-term HIV replication. Increased CD3^+^T cell levels engraftment is obtained, constituted of around 61% of human CD3^+^ thymocytes, 8% of human monocytes in the bone marrow, and 3% of human myeloid DCs in the spleen, 22 weeks after engraftment of mice (48). Since IL-1 β is mainly produced by macrophages and macrophage-like cells, it is tempting to speculate that less protein is secreted after HIV infection in this model as compared to other animal models. In this context, a clear correlation was however achieved between the level of circulating IL-18 and HIV-1 pathogenesis (viral load and loss of CD4^+^ T cells, Figure 3). In summary, all these data demonstrated that the humanized mouse model of HIV infection recapitulates the mean features of inflamasomme activation and is well adapted to assess anti-inflammasome therapies against HIV-1 induced immune activation.

To explore the potential effects of direct inflammasome inhibition *in vivo*, we next evaluated the caspase-1 inhibitor VX-765 administrated as early as day 2 post-infection and for 21 consecutive days. Daily administration of VX-765 significantly reduced circulating levels of IL-18 and levels of circulating pro-inflammatory cytokines, preserved CD4+ T cell homeostasis, and lowered plasmatic viral load and HIV-1 reservoirs in human CD45^+^ immune cells from the spleen. The level of Il-1β was also reduced in the group of VX-765 treated mice as compared to the control group of HIV-infected mice but this difference did not reach statistical significance probably due to a low basal plasma concentration of the cytokine. Interestingly, since a low CD4/CD8 T cell ratio is associated with disease progression to AIDS, our results suggest that VX-765 treatment may delay disease progression and preserve CD4^+^ T cell homeostasis. Pyroptotic CD4^+^ T cells in lymphoid organs was shown through *in vitro* models to release pro-inflammatory cytokines, able to induce a local inflammation that further recruits new CD4^+^ T cells to the site of infection. In addition, innate immune cells recruitment and the activation of an NLRP3 inflammasome might enhance the release of pro-inflammatory mediators. This globally enhances HIV-1 replication, thus stimulating the replenishment of the reservoir. It is worthy to note that treatment with VX-765 reduced significantly plasma viral load and the formation of the HIV-1 reservoir in the spleen of humanized mice. This result is of utmost importance in the modulation of the size of the reservoirs towards HIV-1 cure. Several studies have shown that the timing of cART initiation after infection is crucially related to the size of the reservoir and is decisive in the likelihood of controlling viral replication after analytical treatment interruption (64). Combined early cART treatment with inflamasomme inhibition, if possible during acute HIV infection, might therefore reduce the reservoir’s refill, and should be considered as a therapeutic option to test in HIV-infected patients initiating cART therapy.

We show here that inflammasome activation *in vivo* rather takes places in splenic innate immune CD11c^+^ and CD14^+^ cells than in splenic adaptive CD4^+^ or CD8^+^ or bone marrow immune cells indicating that HIV-1 induced inflammasome activation in splenic CD11c^+^ cells may be the principal source of circulating IL-18. When investigating pyroptosis in subsets of splenic caspase-1 activated cells, we did not find any significant dead cells in CD4^+^ and CD8^+^ T cells. In addition, we did not detect any effect of VX-765 in caspase-1 activation suggesting that the small caspase-1 activation observed in these subsets might reflect a background activity and was not induced by HIV-1 infection. Our *in vivo* data did not confirm previous results obtained *in vitro* by Doitcher et al (82) who proposed pyroptosis as the main mechanism leading to CD4^+^ T cell depletion after HIV-1 infection. We can not exclude that the assay is not sensitive enough in primary human T cells from humanized mice to show pyroptosis or that we did not catch the best time point. HIV-1 infection is associated with programmed cell death, and NLRP3 inflammasome mediated immune T cell depletion was nevertheless shown here by different correlations with the dowstream pathways of NLRP3. Through serial necropsies, Barouch et al. demonstrated that following SIV infection early host response at the mucosal portal of entry and early sites of distal virus spread involved a robust expression of components of the inflammasome and TGF-β pathways which impair innate and adaptative immunity (43). Other anti-inflammatory strategies aiming at breaking this vicious circle and reducing HIV persistence are currently investigated in several clinical trials, including the anti-inflammatory, anti-fibrotic angiotensin II blocker losartan (phase 2, <u>NCT01852942</u>) and angiotensin II receptor antagonist telmisartan (phase 1, <u>NCT02170246</u>), immunosuppressors with mTOR inhibitory activities such as sirolimus (phase 2, <u>NCT02440789</u>) and everolimus (phase 4, <u>NCT02429869</u>) or the anti-inflammatory janus kinase inhibitor ruxolitinib (phase 2, <u>NCT02475655</u>). Most of these strategies target inflammation during chronic HIV infection. HIV-1 induced massive systemic immune activation was repeatidly linked to poor outcomes during chronic infection such as the increased risk of cardiovascular complications (65, 66). Furthermore levels of cytokines IP-10 and IL-18 remain elevated in cART patients compared to healthy donors and elite controllers (67). Therefore complementary approaches are needed to reduce persisting immune activation under cART; inhibiting early inflamasomme activation might be a very promising approach that could be combined to cART for treating acute HIV infection. During chronic HIV-1 infection under cART therapy, residual viral replication and viral protein expression from defective provirusses is a continuous driver of inflammation and might also trigger inflammasome activation (16-18). Furthermore, the incapacity of cART to completely restore immune functions and control chronic inflammation promotes immune aging and the emergence of non-AIDS related diseases such as cancer, cardiovascular diseases, viral infections and cognitive decline (57). Although cART reduce the aberrant cytokine production of monocytes, some inflammatory monocytes subsist while phagocytic monocytes decrease, resulting in a lower antigen presentation to CD8^+^ T cells (68). Taken together, all these data suggest that inflamasomme inhibition by offering simultaneous reduction of inflammation and HIV-1 reservoir might be evaluated to support cure research.

Besides HIV-1, a number of viruses such as the influenza virus or measles virus have evolved mechanisms to suppress the NLRP3 inflammasome (69). More recently, it has been proposed that inhibiting the inflammasomes pathways and/or pyroptosis in COVID-19 therapy would be an efficient strategy to decrease SARS-CoV-2 virulence and NLRP3-mediated inflammation for treatment of severe COVID-19 disease (70). NLRP3 modulation might therefore be a promising intervention to balance a functional immune response for infection control of different viruses. Novel mechanisms for NLPR3 modulation have been proposed to promote an effective innate immune response against fungal pathogens (71) highlighting that further investigations need to be conducted to understand the adapted control of NLRP3. In conclusion, our results demonstrate the benefits of inhibiting inflammasome early after HIV-1 infection and suggest to combine such therapies to current early cART. Since administration of cART will be less intrusive with long acting formulations and combined with other therapies in the context of HIV cure in the near future, we strongly support the evaluation of inflammasome inhibitors in HIV-infected patients.

---

**Figure Suppl 1.**
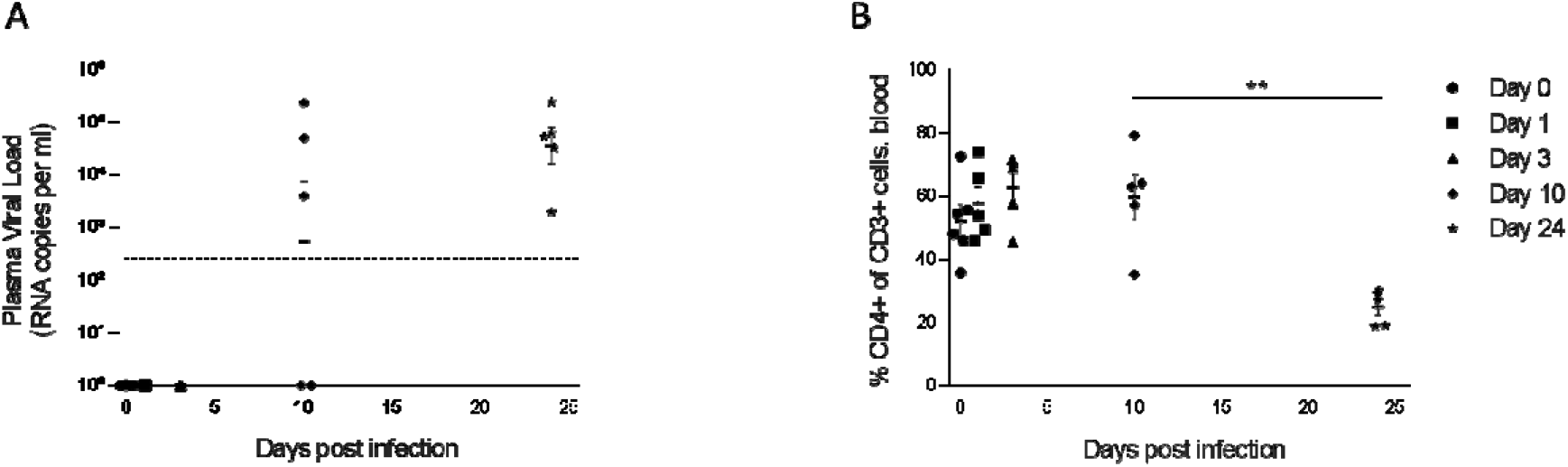
Kinetics of viremia and early CD4^+^ T cell loss during HIV-1 infection of huNSG mice. (A) Plasma HIV-1 viral load measured by ddPCR (LOD: 235 copies/ml) and (B) human CD4^+^ T cells percentage among CD3^+^ cells measured by flow in HIV-1_JRCF_ i.p. infected NSG humanized mice at day 0 (n=6), day 1 (n=5), day 3 (n=5), day 10 (n=5) and day 24 (n=5). Statistical tests were analysed by Mann–Whitney t-tests for comparison of two groups.

**Figure Suppl 2.**
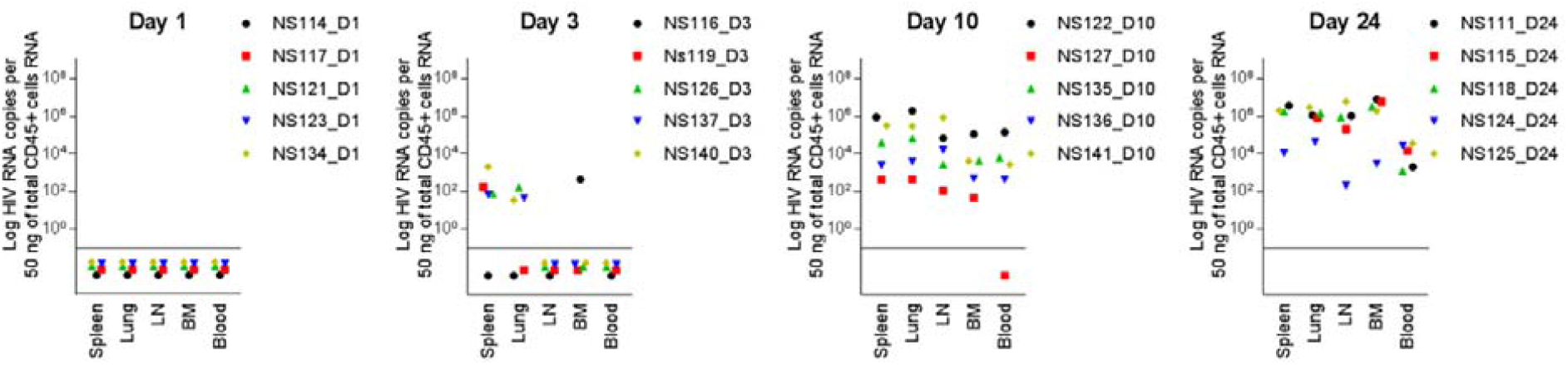
HIV-1 disseminates within 3 days post infection in huNSG mice. Viral RNA in the spleen, lung, lymph nodes (LN), bone marrow (BM) and in the blood were measured by RT-PCR (Log HIV RNA copies per 50 ng of total hCD45^+^ cells RNA) in HIV-1_JRCF_ i.p. infected NSG humanized mice at day 1 (n=6), day 3 (d=5), day 10 (n=5) and day 24 (n=5). Samples in which viral RNA was below 40 copies per 50 ng of total CD45^+^ cells RNA were plotted below the horizontal line.

**Figure Suppl 3.**
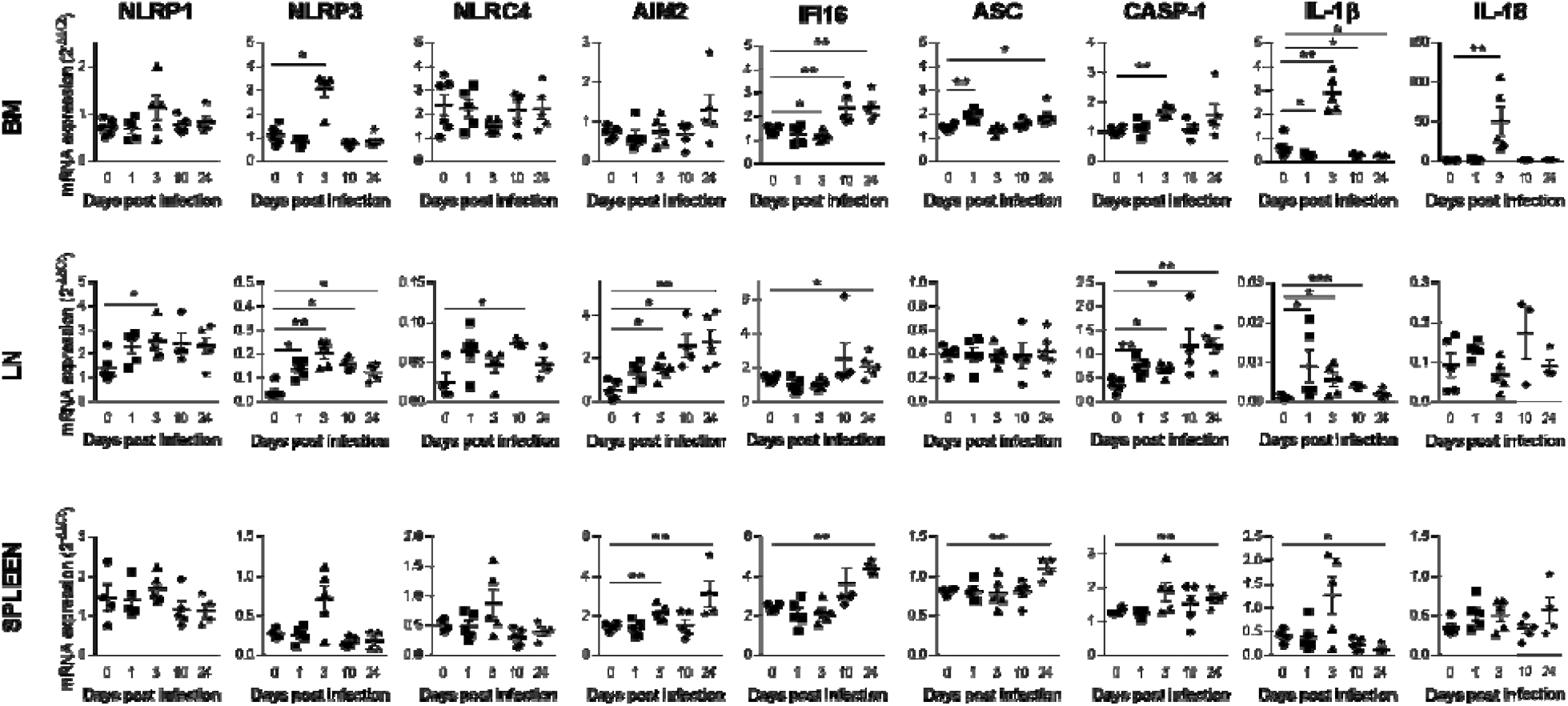
HIV-1 induction of inflammasome related genes expression in tissues. Relative mRNA expression of the indicated genes measured by qPCR in hCD45^+^ cells isolated from bone marrow (BM), lymph nodes (LN) and spleen in HIV-1_JRCF_ i.p. infected huNSG mice at day 0 (n=5), day 1 (n=5), day 3 (n=5), day 10 (n=6) and day 24 (n=5). Results were analyzed using the 2^-ΔΔCT^ method. Statistical tests were analysed by Mann–Whitney t-tests for comparison of two groups

**Suppl. Figure 4.**
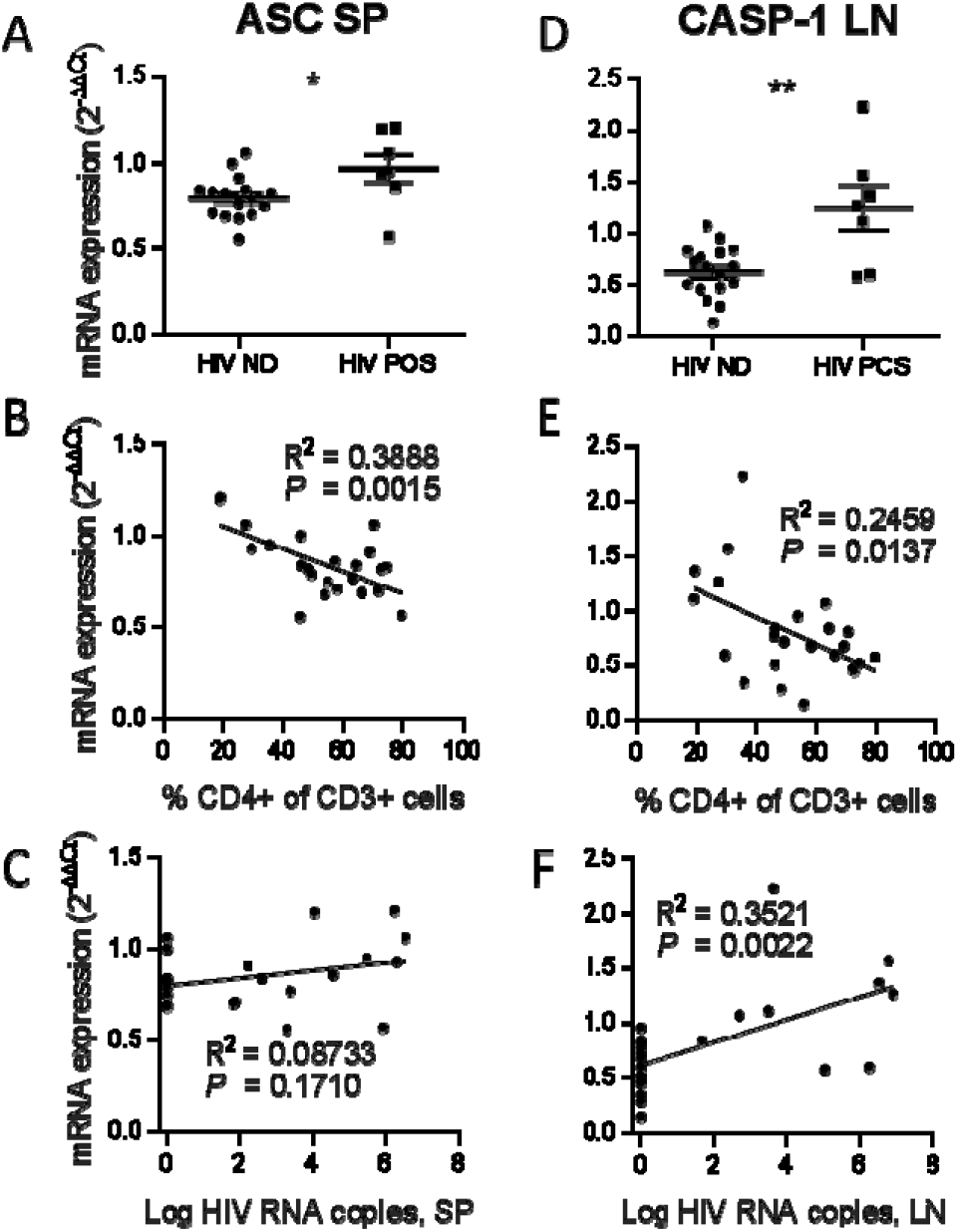
ASC and CASP-1 expression correlates with HIV-1 replication and pathogenesis. ASC and CASP-1 relative mRNA expression comparison between viremic (HIV POS) and aviremic (HIV ND) huNSG mice (A and D) in the spleen (SP) and in the lymph nodes (LN). Correlation between ASC and CASP-1 relative mRNA expression with corresponding tissue viral RNA copies (B and E) or CD4^+^T cells percentage in the blood (C and F). qPCR results were analyzed using the 2^-ΔΔCT^ method. Statistical tests were analysed by Mann–Whitney t-tests for comparison of two groups or Spearman correlation tests.

**Suppl. Figure 5.**
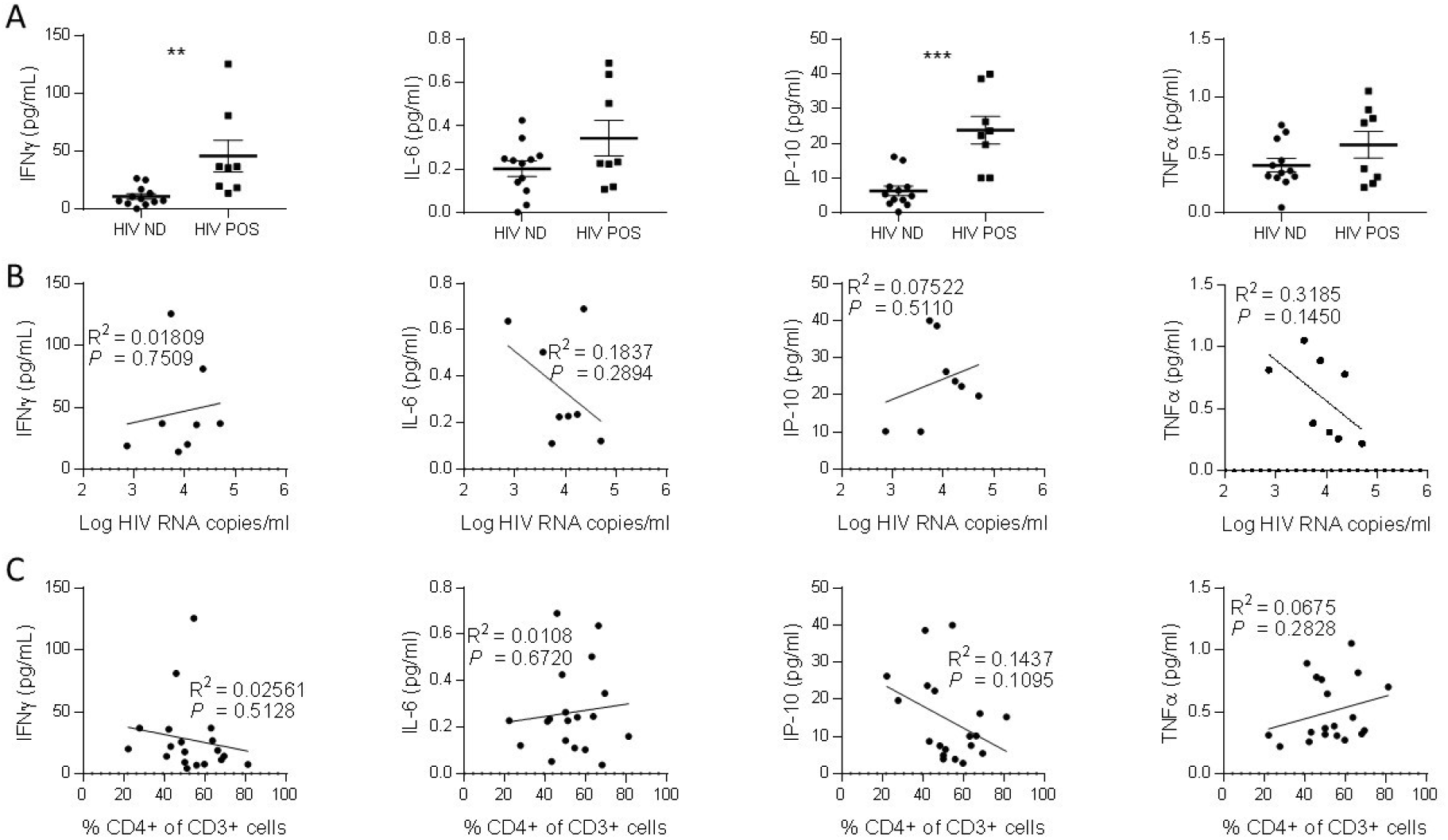
IFN-γ and IP-10 are inducted by HIV-1 replication. (A) Plasmatic cytokines levels comparison between viremic (HIV POS) and aviremic (HIV ND) huNSG mice. Correlations between plasmatic cytokines levels with the plasmatic viral load (B) or CD4^**+**^ T cells percentage in the blood (C) at day 1 (n=5), day 3 (n=5), day 10 (n=6) and day 24 (n=5). Statistical tests were analysed by Mann–Whitney t-tests for comparison of two groups.

**Suppl. Figure 6.**
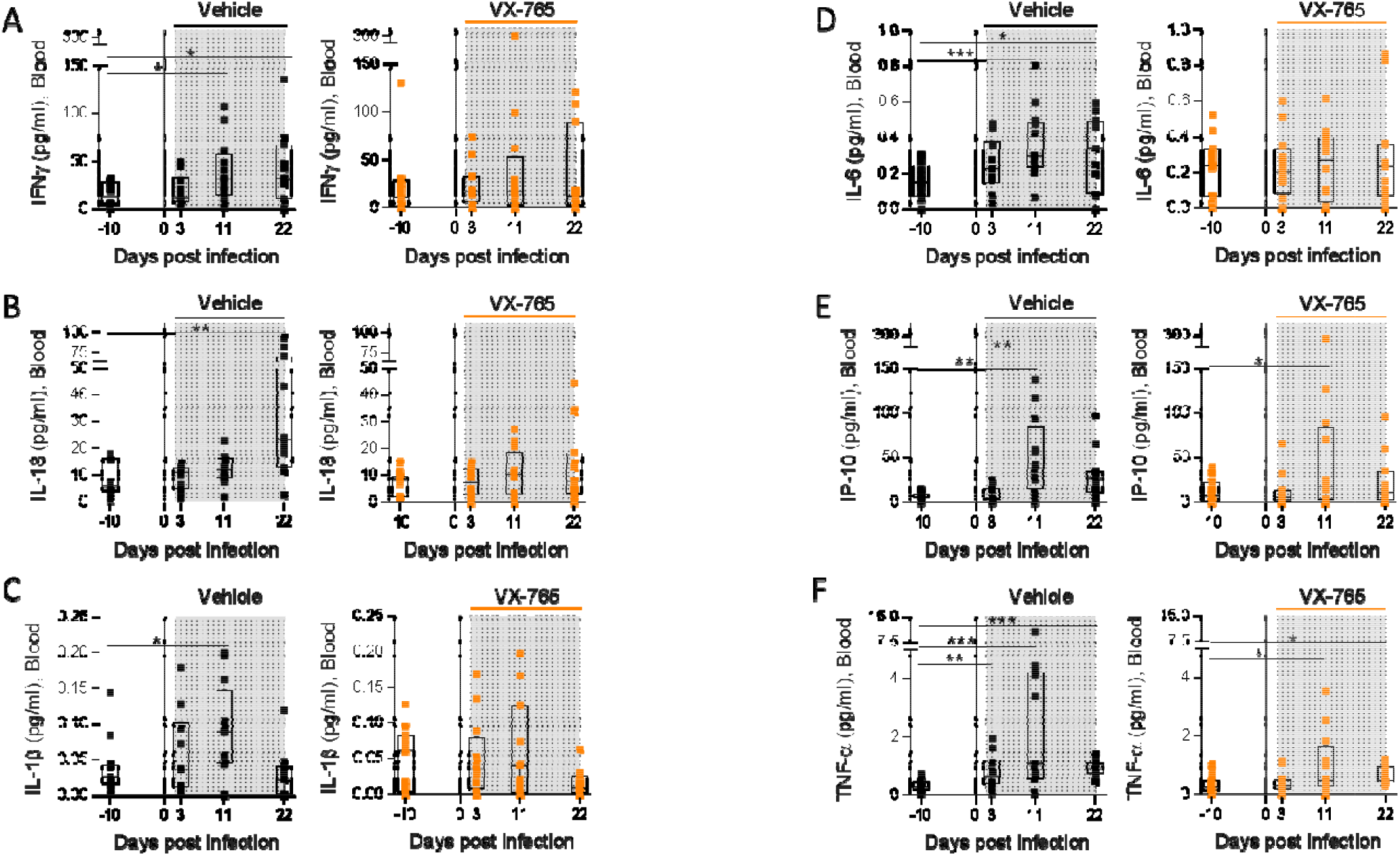
HIV-1 infection increases pro-inflammatory cytokines circulating levels. Plasmatic cytokines concentration measured using a multiplex U-PLEX Biomarker assay kit in longitudinal plasma samples of HIV-1_JRCF_ i.p. infected huNSG mice treated with VX-765 (VX-765, n=12) or vehicle (Vehicle, n=12). Statistical tests were analysed by Wilcoxon’s matched pairs signed rank test.

**Suppl. Figure 7.**
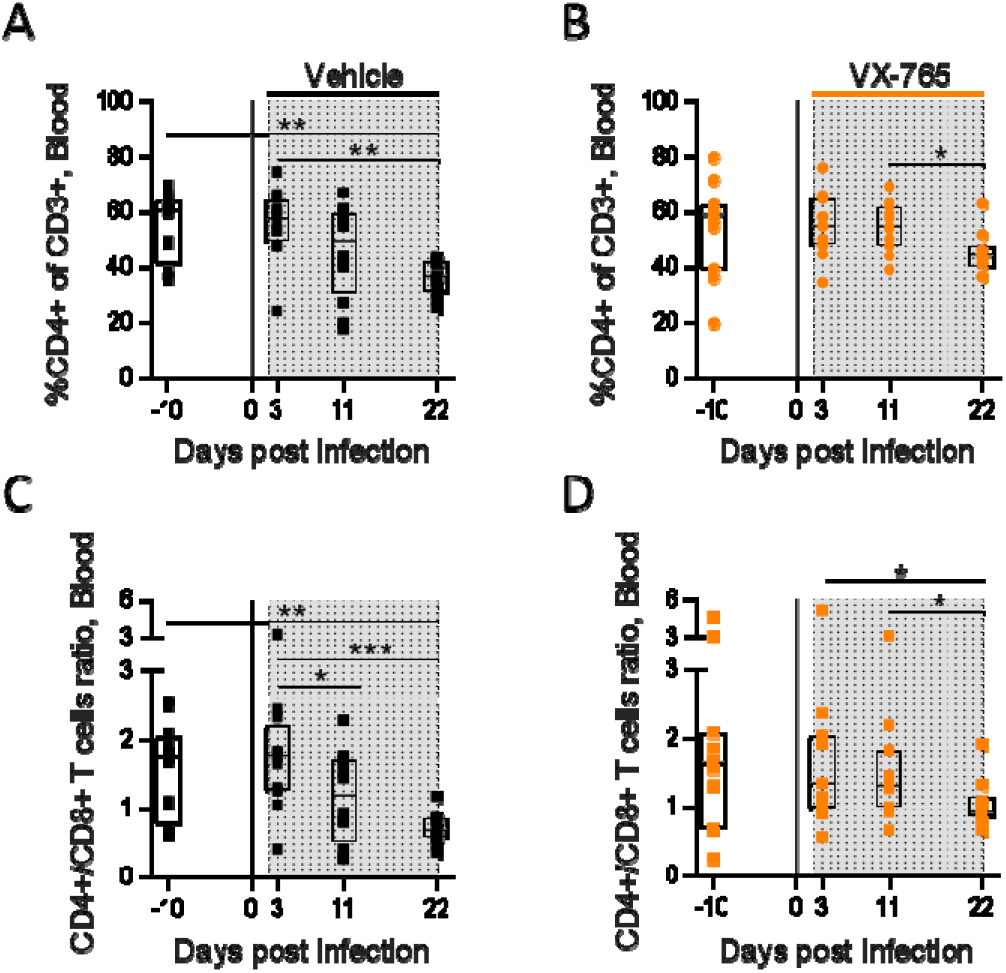
VX-765 treatment prevents CD4^+^ T cell depletion in the blood. Blood human CD4^+^ T cells percentage among CD3^+^ cells (A and B) and CD4^+^/CD8^+^ T cell ratios (C and D) measured longitudinally by flow cytometry in vehicle or VX-765 treated HIV-1 infected HuNSG mice. Statistical tests were analysed by Wilcoxon’s matched pairs signed rank test.

**Suppl. Figure 8.**
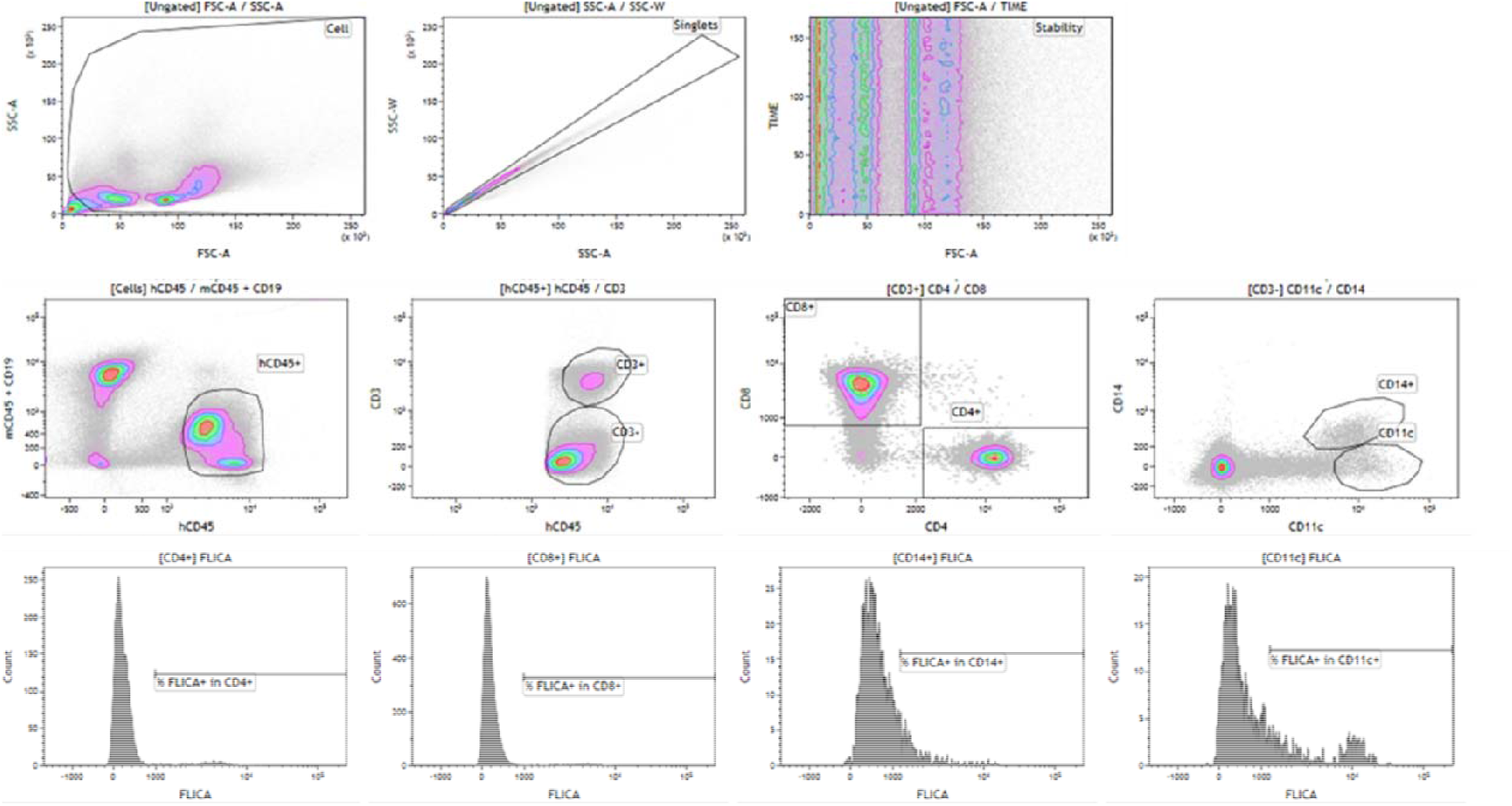
Gating strategy for FLICA analysis.

**Suppl. Figure 9.**
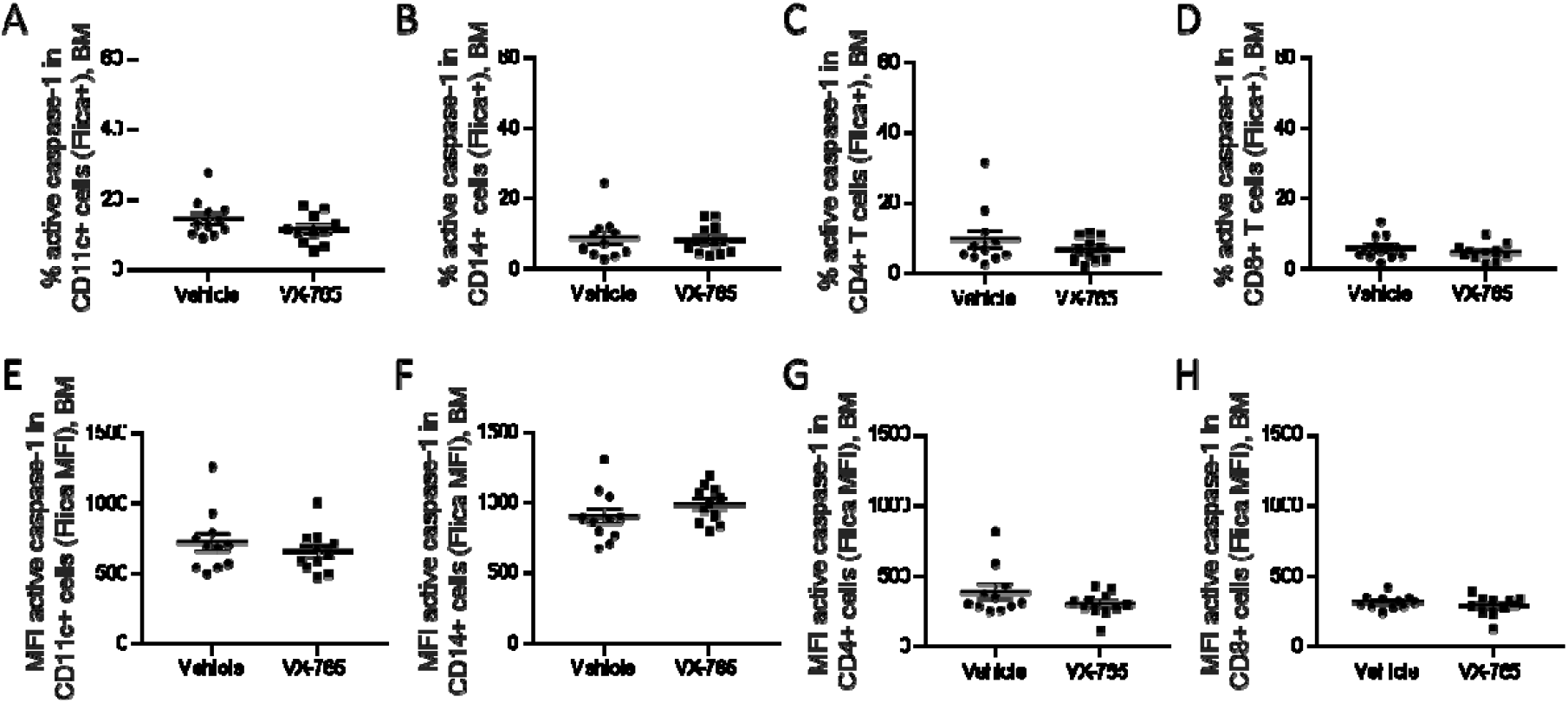
VX-765 treatment does not affect caspase-1 activation in the bone marrow. Cells from the bone marrow were stained for active caspase-1 using FAM-FLICA reagent. Percentage of FAM-FLICA^+^ CD11c^+^ (A), CD14^+^ (B), CD4^+^ (C), CD8^+^ (D) splenic cells in vehicle or VX-765 treated HIV-1 infected HuNSG mice at day 22 post-infection. Mean fluorescence intensity (MFI) of FAM-FLICA in CD11c^+^ (E), CD14^+^ (F), CD4^+^ (G), CD8^+^ (H) splenic cells in vehicle or VX-765 treated HIV-1 infected HuNSG mice at day 22 post-infection. Statistical tests were analysed by Mann–Whitney t-tests for comparison of two groups.

**Suppl. Figure 10.**
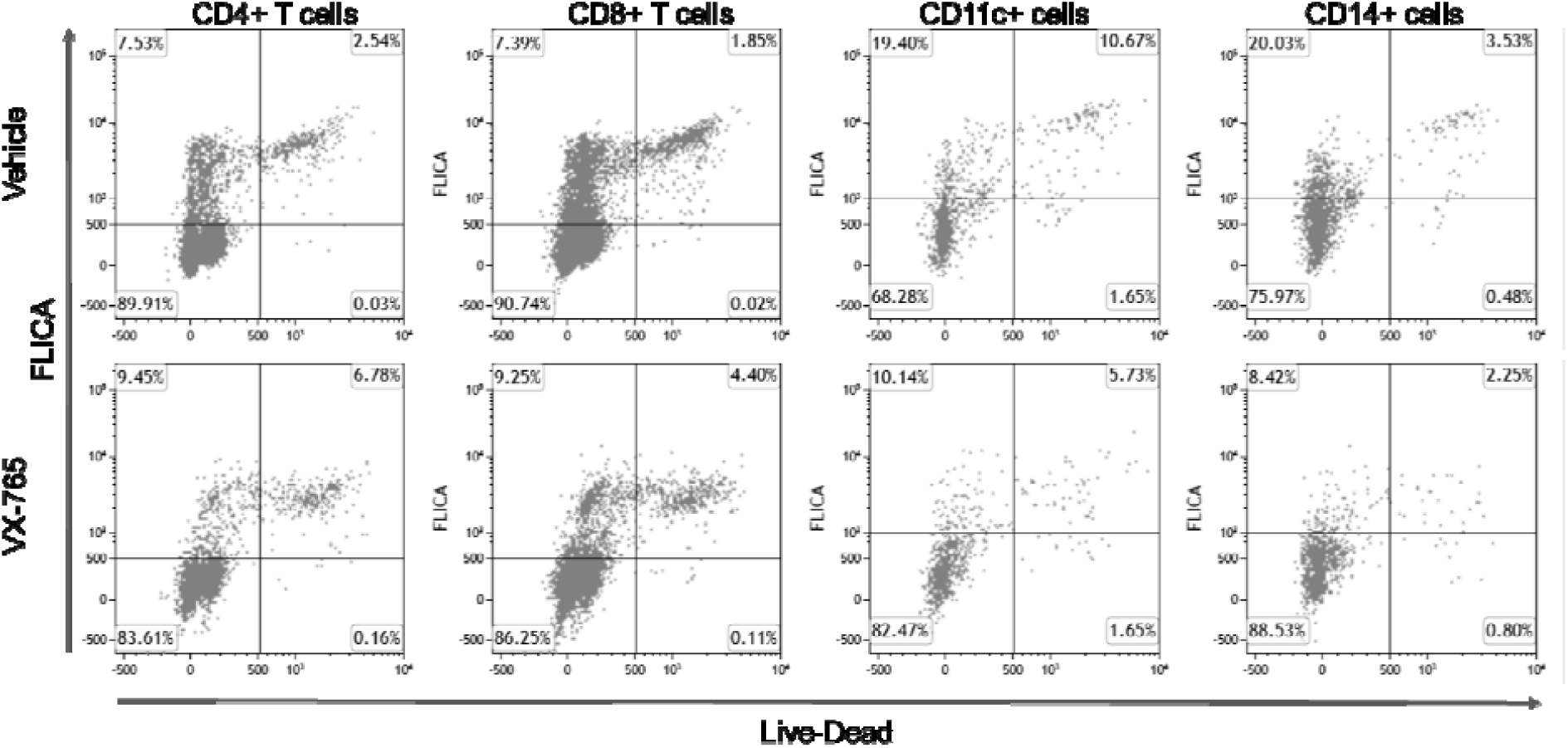
Representative dot plots of FLICA vesrsus Live-dead staining in splenocytes from VX-765 and vehicle treated HIV-1 infected HuNSG mice.

